# SSR42 is a Novel Regulator of Cytolytic Activity in *Staphylococcus aureus*

**DOI:** 10.1101/2024.07.11.603084

**Authors:** Mary-Elizabeth Jobson, Brooke R. Tomlinson, Emilee M. Mustor, Emily A. Felton, Andy Weiss, Clayton C. Caswell, Lindsey N. Shaw

**Author notes:** Address correspondence to Lindsey N. Shaw, PhD.

## Abstract

SSR42 is the longest noncoding RNA in the *S. aureus* cell and the second-most abundant transcript in the stationary phase transcriptome, second only to RNAIII. It is highly conserved across strains and exhibits pronounced stability in stationary phase, however the mechanism behind its regulatory role has yet to be fully elucidated. Herein, we used transcriptomic and proteomic approaches to probe the role of SSR42, revealing that it is a powerful, novel activator of the primary leukocidin LukAB. SSR42 is required for cytotoxicity towards, and escape from within, human neutrophils, and also mediates survival within human blood. We show that SSR42 wields this role via derepression by the peroxide repressor PerR in response to the presence of human neutrophils and governs *lukAB* induction in this niche. Importantly, this regulation is driven by direct RNA-RNA interaction, as we show binding of the 5’ UTR of the *lukAB* transcript with the 3’ end of SSR42, which ultimately modulates transcript stability as well as translational activity. Finally, we demonstrate that this behavior is absolutely required for full virulence of *S. aureus* in murine models of both pneumonia and sepsis. Collectively, we present SSR42 as a pleiotropic regulatory RNA that acts as a nexus between environmental sensing and the regulation of pathogenesis, responding to environmental stimuli and host immune factors to bolster cytotoxic behavior and facilitate infection in *S. aureus*.

**Importance:** *S. aureus* is a master pathogen due to its formidable collection of virulence factors. These are tightly controlled by a diverse group of regulators that titrate their abundance to adapt to unique infectious niches. The role of regulatory RNAs in stress adaptation and pathogenesis is becoming increasingly more relevant in *S. aureus*. In this study, we provide the most comprehensive global analysis to date of just such a factor, SSR42. Specifically, we uncover that SSR42 is required for mediating cytotoxicity - one of the pillars of infection - in response to phagocytosis by human neutrophils. We find that SSR42 is induced by components of the host immune system and facilitates downstream activation of cytotoxic factors via RNA-RNA interactions. This illustrates that SSR42 forms a pivotal link between sensing the external environment and mediating resistance to oxidative stress while promoting virulence, solidifying it as a major global regulator in *S. aureus*.

## Introduction

Over the last few decades, more than 300 regulatory RNAs have been identified in *Staphylococcus aureus* strains (1–6). While progress has been made exploring these elements mechanistically, there is still a dearth of knowledge on the role of regulatory RNAs in complex regulatory systems, as compared to classical transcription factors. Commonly referred to as small RNAs (sRNAs), these factors range from *trans*-encoded-to antisense-transcripts, typically acting in a posttranscriptional manner to allow for rapid adaptation to environmental cues (6–13). The most well-studied example of an sRNA in *S. aureus* is RNAIII, which is the effector for the Agr system. It is known to modulate mRNA abundance by relieving inhibitory secondary structures at the Shine-Dalgarno sequence (e.g. *hla*), or by occluding this site to block access of the ribosome (e.g. *spa, rot, sbi*) ultimately promoting transcript degradation (14–17).

Many other virulence-related regulatory RNAs have also been documented in *S. aureus*, such as the Teg family of transcripts, which include Teg41 that impacts hemolytic ability through engagement with the PSMs, and Teg58 which monitors arginine dysbiosis and modulates biofilm production (18–20). The Spr-group of regulators form another family of regulatory RNAs, and includes SprC, which plays a role in controlling metabolism and immune evasion, SprY, which works to titrate RNAIII activity through direct ‘sponge’ interactions, and SprD, which works in consort with RNAIII to repress *sbi* (21–24) . RsaC is a ∼1,100-nt transcript that is expressed during manganese starvation and enhances the oxidative stress response, leading to increased survival during confrontation with the host immune system; thus forming a link between metal homeostasis and pathogenesis (25). From a broader perspective, a recent global study utilizing CLASH technology to uncover RNA-RNA interactions in host-like conditions identified hundreds of binding events, occurring not only between regulatory RNAs and mRNAs, but between regulatory RNAs themselves as well. Some of these interactions linked the metabolic status of the cell to the expression of membrane-permeabilizing toxins, while furthermore uncovering an RNA-sponge relationship between RsaE and RsaI (13)

The largest regulatory RNA in *S. aureus* is SSR42, which is 1,233 nt long, contains no functional ORFs, and bears 98% sequence conservation across strains, with only two single nucleotide variants found in UAMS-1 (26). It was first characterized by Morrison et al. from a larger group of <u>S</u>mall <u>S</u>table <u>R</u>NAs (SSRs), where it was shown that the SSR42 transcript exhibits enhanced stability during stationary phase growth, with a half-life of over 30 minutes, compared to a ∼5 minutes in exponential phase. Using microarray analysis, 80 mRNA species were found to be altered in an SSR42 mutant strain, with key virulence factors such as *hla, lukF, hlgC, aur,* and *sraP* found to be differentially expressed. Phenotypically, an SSR42 mutant demonstrated an abrogated ability to lyse human erythrocytes and impaired virulence in a murine model of skin abscess formation (26). Others have shown that SSR42 is tightly controlled via the Repressor of Surface Proteins (Rsp), and in turn may control expression of the two-component system SaeRS, potentially explaining the ahemolytic nature of SSR42 mutant strains (27). Work by our group identified that SSR42 is unique to *S. aureus* and does not exist in other Staphylococcal species (5). We also demonstrate that expression of SSR42 is profoundly increased during the growth of *S. aureus* in human serum (1).

Despite SSR42 being well linked to *S. aureus* pathogenesis, no direct target or mechanism of action has been demonstrated for this molecule. Herein, we use a combination of large-scale omics studies coupled with genetic and biochemical analyses to demonstrate that SSR42 is a novel regulator of *lukAB* expression and cytolytic activity through direct RNA-RNA interaction. We show that this regulation is controlled by the peroxide sensor PerR and is stimulated in response to human neutrophils, mediating escape from the intracellular niche. Finally, the impact of this interaction is illustrated by impaired virulence in murine models of both sepsis and pneumonia.

## Methods

### Toxicity towards neutrophils

Cytotoxicity assays were performed as previously described (28), with some variations. The immortalized HL-60 cell line was used as a model of human neutrophil engagement. Cells were grown to confluence in RPMI supplemented with 10% FBS and 100U Pen/Strep in 37°C and 5% CO_2_. Upon reaching sufficient cell density, cells were differentiated into neutrophil-like cells by the addition of 1.25% DMSO for 4 days; which was confirmed via observation of cell morphology. For use, 1×10^5^ cells were seeded in 100µl RPMI with 10% FBS into a 96-well plate. Bacterial cultures were grown in biological triplicate overnight and standardized to 0.05 before being grown for 15h. Following incubation, cultures were pelleted via centrifugation and supernatant was diluted two-fold in TSB. Seeded neutrophils were intoxicated in technical duplicate with 5µl of diluted bacterial supernatant (5% total volume) followed by incubation at 37°C, 5% CO_2_ for 1h. Cell viability was measured using the CellTiter96 Aqueous One Solution Cell proliferation reagent (Promega). Following 1h incubation, 20µl of the CellTiter reagent was added to each well, and the plate was returned to the incubator for 2h. Color development was assessed via OD_490_ using a Biotek Cytation 5 plate reader. Data is reported as percent viability of supernatant-treated cells with respect to neutrophils treated with 5µl TSB only.

### Intracellular survival within neutrophils

To assess intracellular survival within human neutrophils, HL-60 cells were grown and differentiated as described above, and 2×10^6^ cells were seeded into 500µl in a 24-well plate. Bacterial cultures were also grown and processed as above, with the following modifications. After centrifugation, pelleted cells were washed 2X with PBS, following which all strains were standardized to 10x the desired final MOI of 30 (OD_600_=0.135 for *S. aureus*), thus all strains were standardized to an OD_600_=13.5 in PBS. From these standardized cultures, 5µl was added to each well of neutrophils and incubated for 1h at 37°C and 5% CO_2_. Following incubation, cells were removed from wells and pelleted at 2.5 x *g* for 1 minute before being resuspended in pre-warmed PBS. This PBS wash was repeated, and the cells were finally resuspended in pre-warmed RPMI supplemented with 30µg/mL gentamicin and returned to incubate for 24h. At this point in time the cells were removed and washed twice with pre-warmed PBS once again, however the final resuspension was in 0.5% Triton X-100 to lyse neutrophils. Lysates were serially diluted and plated to enumerate CFU of internalized bacteria. Viability of the infected neutrophils was also quantified after the 24h infection period by adding 100µl of CellTiter reagent to each well, incubating for 1h at 37°C and 5% CO_2_, and measuring OD_490_ using a Biotek Cytation 5 plate reader as described above.

### Measuring translational control by SSR42 via GFP reporters

SSR42 mutant strains containing both the *lukAB* mRNA-GFP fusion plasmid (pCN33) alongside the SSR42 expression or empty vector plasmid (pICS) were grown overnight in biological triplicate. Cultures were synchronized for 3h and standardized to an OD_600_ of 0.05 in a black-walled, clear bottom 96-well plate. Strains were then monitored for OD_600_ and fluorescence (excitation:485/20, emission:528/20, Gain:75) measured every 15min for 18h. Fluorescence levels of the mRNA reporter in the presence of SSR42 *in trans* were compared to fluorescence levels of the mRNA reporter in the presence of an empty vector. Strains were normalized by growth via OD_600_.

### Electrophoretic mobility shift assays (EMSAs)

All RNA was generated via *in vitro* transcription using a T7 RNA polymerase promoter and the MAXIscript T7 kit (ThermoFisher). SSR42 was amplified using OL7499/OL7500 while fragments of the *lukA* UTR and *splE* UTR were amplified using primers 7501/7502 and 7523/7524, respectively. These T7-controlled fragments were cloned into pGEM-T Easy and liberated from the plasmid using restriction enzymes NcoI and PstI (Thermo Fisher Scientific). *In vitro* transcription was performed using the MAXIscript T7 kit according to manufacturer instructions and using 1µg of template DNA, ^32^P-labeled UTP for the mRNA, and unlabeled UTP for SSR42. TurboDNase was added to degrade any remaining DNA. Reactions were cleaned using a Nucleotide Clean-up Kit (Qiagen). RNA-RNA binding reactions were performed with labeled mRNA and SSR42 at varying concentrations, which were heated to 90°C for 2 minutes and then incubated at RT for 30 min. Native acrylamide gels were equilibrated on ice for 30 minutes in 0.5x TBE before loading binding reactions, which were mixed with RNA loading dye and then run at 100V until the dye front has moved ¾ of the way through the gel. Gels were dried and affixed to filter paper using a gel dryer before being visualized via autoradiography.

### Murine models

All animal studies in this work were performed in accordance with and approved by the Institutional Animal Care and Use Committee of the University of South Florida. To prepare inocula for a model of bacterial sepsis, bacteria was washed 2x with PBS before being adjusted to 5×10^8^ CFU/mL in PBS. Female, six-week-old CD-1 mice (Charles River Laboratories) were inoculated with 100µl of bacteria, resulting in a final dose of 5×10^7^. Infection was allowed to progress for 7 days, or until mice reach a premoribund state, at which point they were euthanized. After the infection period, all mice were euthanized and the brain, heart, liver, spleen, lungs, and kidneys were harvested. For murine models of pneumonia, inocula were prepared in a similar manner, however the bacterial density was instead adjusted to equal a final dose of 1×10^8^ in 30 µl. Female and male 6-week old C57BL/6 mice were intranasally inoculated and infections were allowed to proceed for 24h, after which mice were euthanized and lungs harvested. Organs for both models were homogenized in 2mL PBS and serially diluted and plated for CFU/mL. All infection experiments were performed with 8 mice per strain and a Mann-Whitney test was used to measure statistical significance.

### Data Availability & Additional Methods

RNAseq data from this study is available under the GEO accession #GSE237701. Proteomics data is available via proteomeXchange under the ID PXD043724. Additional methods can be found in the supplemental file.

## Results

### SSR42 is global regulator of virulence factor expression

Although previous studies exist exploring SSR42, the only large-scale omics examination of this factor is a microarray performed over a decade ago (26). Thus, to bring new insight to the role of SSR42, we created a complete SSR42 deletion in the USA300 background and performed RNA-sequencing of the WT and mutant strains. To confirm validity of our mutant strain, an analysis of RNAseq reads revealed the expected deletion of SSR42 **(Fig. S1A),** with concomitant downregulation of this gene observed (−2,955-fold compared to WT). We also determined that this deletion had no impact on growth kinetics **(Fig. S1B).** When assessing reads from the wild-type dataset, we noted that SSR42 was the second most abundant transcript in the cell, second only to RNAIII **(Fig. S2).** Moreover, 20% of all RNA in stationary phase cells under these conditions (absent rRNA) belongs to SSR42, and 45% to SSR42 and RNAIII combined. Beyond this, we found a total of 220 genes were significantly differentially expressed between wild-type and ΔSSR42 (**Fig. 1A)**, greatly surpassing the 80 genes previously shown to be controlled by SSR42 via microarray. These included those involved in metabolism, transport, signal transduction and, by far the largest functional category (33 such factors), human disease. Regarding this latter group, SSR42 primarily serves as an activator, with 83% of transcripts decreased in abundance in the mutant strain **(Fig. 1BC).**

**Figure 1.**
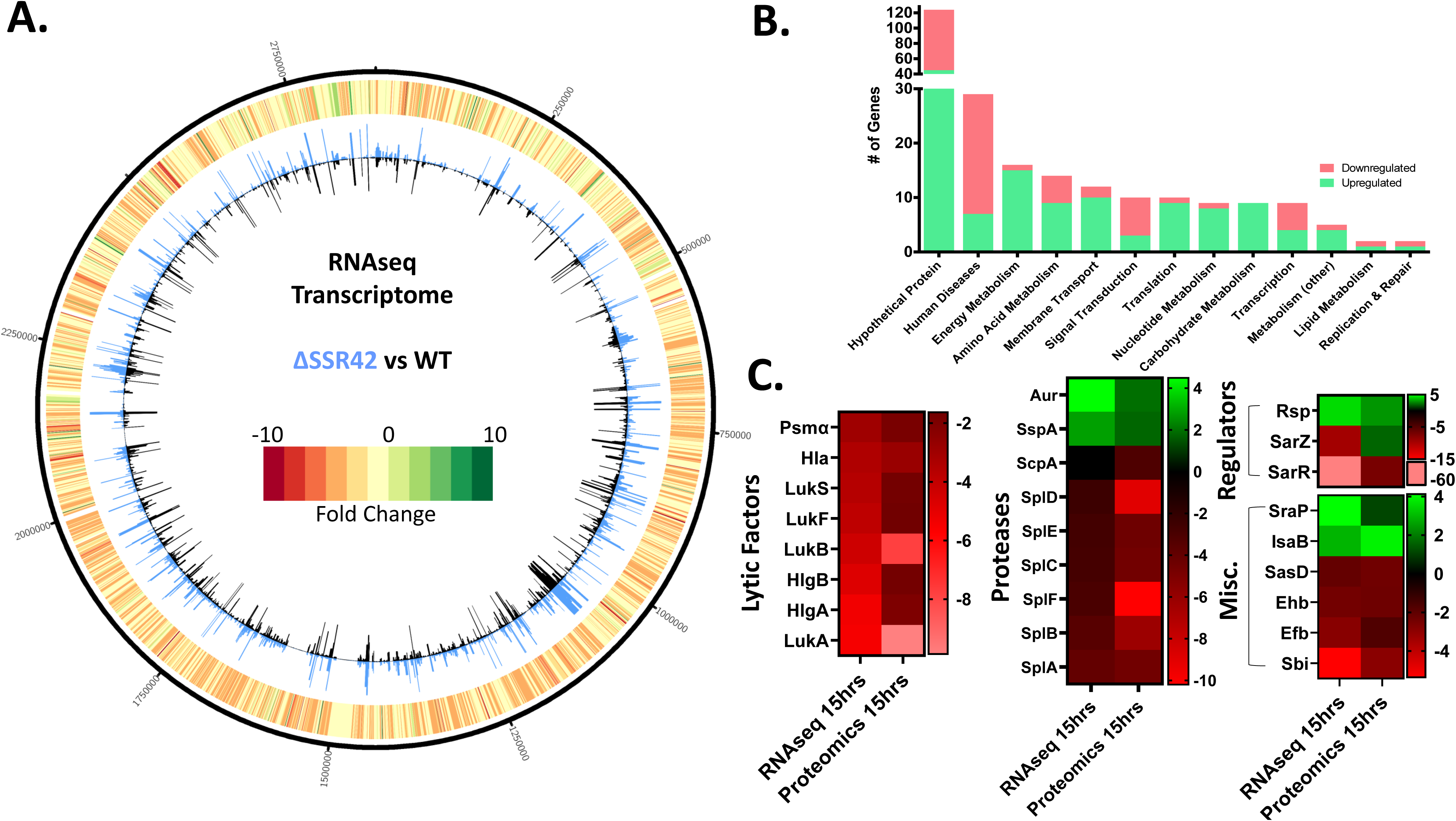
SSR42 is a global regulator of virulence factor production. **(A)** CIRCOS file illustrating transcriptional changes between wild-type and ΔSSR42 at 15h of growth. The inner circle exhibits transcript expression represented as a histogram of TPM expression values for the wild type (black bars) and SSR42 mutant (blue bars). Outermost circle is a heatmap describing fold change in expression where green indicates a positive fold change and red indicates a negative fold change in comparison to the wild type. **(B)** The number of genes upregulated (green) or downregulated (pink) ≥2-fold in the SSR42 null strain at 15h, sorted by function based on KEGG ontology. **(C)** Heat map illustrating changes in transcript and protein abundance of key virulence factors of interest.

Key virulence factors regulated by SSR42 included secreted toxins, such as alpha hemolysin (*hla,* -3.44-fold in ΔSSR42 versus WT), gamma hemolysin (*hlgA* -5.39, *hlgB* - 4.77), the Panton-Valentine leukocidin (*lukS* -3.62, *lukF* -3.86), and the *lukAB* leukocidin (−5.49, -4.29). Secreted proteases were also impacted, with all serine protease (*spl*) operon genes downregulated in the mutant (∼-3-fold), whilst aureolysin (*aur*, +4.5) and the V8 protease (*sspA*, +2.81) were both increased in abundance. Moreover, SSR42 appears to activate immune evasion factors, including igG-binding protein *sbi* (−5.36), extracellular fibrinogen-binding protein *efb* (−2.85), and staphylokinase *sak* (−2.48). In the same vein, we identified a profound change in the secreted extracellular matrix and plasma binding protein *emp* (−34.6). Other virulence factors increased in abundance in ΔSSR42 include the serine-rich adhesin for platelets *sraP* (+4.12) and the Immunodominant Antigen B *isaB* (+2.92).

We also uncovered alterations in known transcriptional regulators, the most striking being the SarA-family proteins *sarZ* (−9.6) and *sarR* (−62.3), which were downregulated in ΔSSR42 compared to wild-type. Interestingly, the Repressor of Surface Proteins (*rsp*), which is divergently transcribed from SSR42 and has been shown to be a primary activator of its expression (27), was found to be upregulated 4.31-fold upon SSR42 deletion. Finally, we identified changes in expression of the metalloregulators *fur* (+3.14) and *mntR* (−2.13), as well as the alternative sigma factor *sigS* (−2.75), in the mutant strain. A random subset of these fold changes were explored and confirmed by qPCR analysis (**Fig S3A**).

We next used proteomic analysis to validate these changes, revealing strong correlation to our RNAseq data **(Fig. 1 A-C)**. Notably, while leukocidins LukSF and LukAB exhibit decreased protein abundance in the mutant that matches RNAseq trends, LukAB in particular exhibits an even more profound decrease in fold-changes at the protein level (RNAseq: *lukA* = -5.51, *lukB* = -4.3; Proteomics - LukA = -9.87, LukB = -7.83). A complete list of all fold changes for both omic studies can be found in **Table S1** and a comparison between the two datasets can be found in **Fig. S3B.**

### SSR42 is a novel regulator of cytolytic activity

Given that LukAB levels had such profound changes in our -omic studies, we next set out to explore if this had a physiological relevant impact on leukocidin activity. As LukAB is the primary effector for the lysis of human neutrophils (29), human promyeloblast (HL-60) cells were differentiated into neutrophils before being treated with bacterial supernatant from the wild type, mutant, and complemented strains. Strikingly, after a 1h incubation, supernatant from ΔSSR42 elicited markedly reduced killing compared to the wild-type and complement **(Fig. 2A).** To see how this translated into bacterial viability, we next explored how deletion of SSR42 impacted *S. aureus* survival during exposure to whole human blood. Here we noted the mutant exhibited a 7-fold increase in survival compared to the wild type and complementing strains **(Fig. 2B).** Of note, *lukAB* mutants of *S. aureus* have been shown to have increased intracellular abundance within neutrophils (a key component of whole blood), due to an inability to use the toxin to escape the phagosome (30). To determine if this was the case for ΔSSR42, we next measured intracellular loads of our strains within HL-60 cells. Here we determined that, after 24h incubation, the mutant displayed a dramatic increase in intracellular abundance compared to the wild type and complement within this niche **(Fig. 2C).** We also measured the viability of infected neutrophils following the 24h infection period, revealing increased viability in cells infected with ΔSSR42 in comparison to the wild-type; likely due to decreased cytotoxicity of the mutant strain **(Fig. S4).** Although this trend is not statistically significant, complementation of SSR42 on a multi-copy plasmid causes a profound decrease in cell viability in comparison to ΔSSR42. As such, we conclude that SSR42 is a major new regulator of LukAB activity in *S. aureus*, mediating exoenzyme-based cytotoxicity as well as intracellular escape from the phagosome.

**Figure 2.**
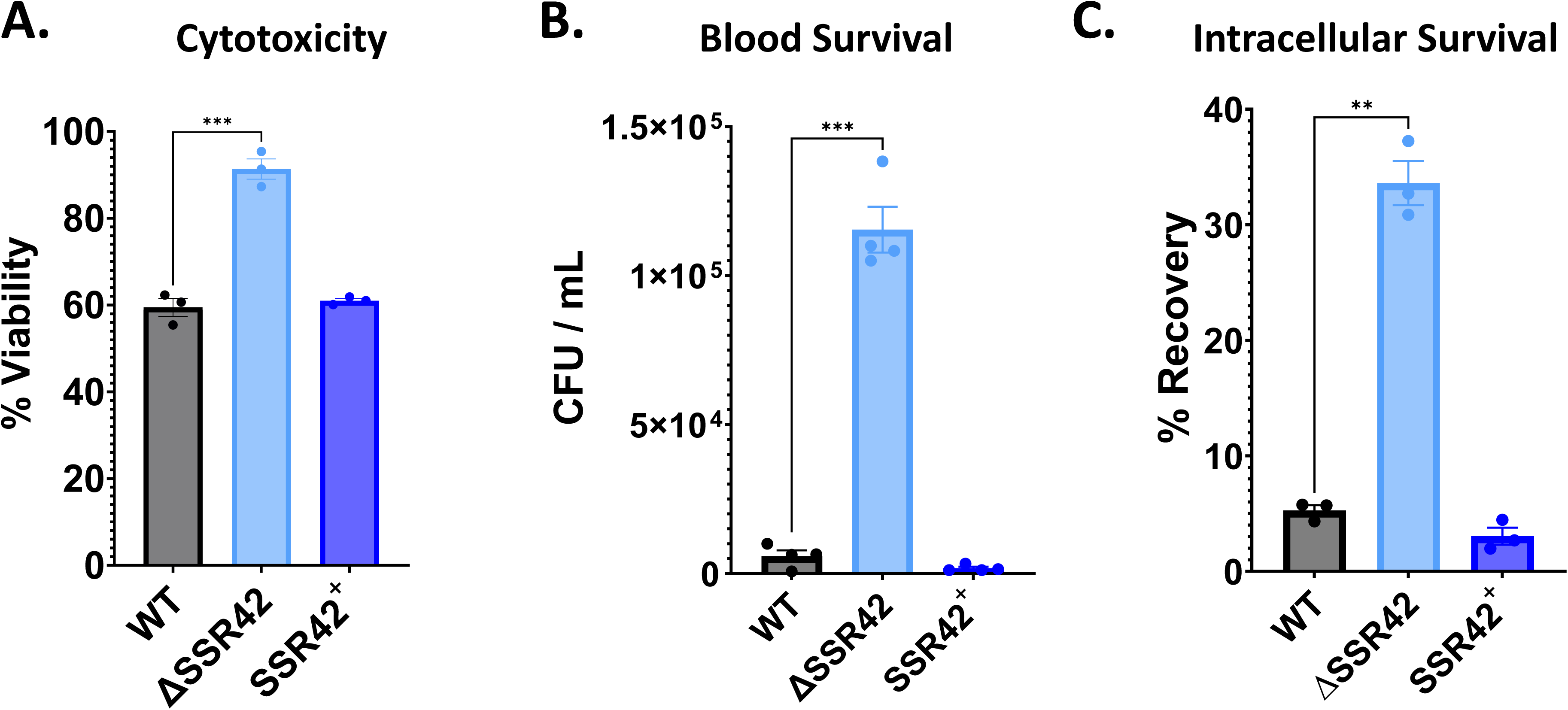
SSR42 plays a central role during engagement with the innate immune system. **(A)** HL-60 cells were differentiated into neutrophil-like cells with 1.25% DMSO for 4 days before being treated with 5% v/v bacterial supernatant from 15 h cultures of the strains shown. Intoxicated cells were incubated for 1 h before cell viability was measured using the CellTiter reagent. Percent cell viability in comparison to TSB-treated cells is shown. **(B)** Bacterial strains were synchronized and then standardized to an OD_600_ of 0.05 in 1 mL pooled-donor, whole human blood. Bacterial recovery after 4 h was enumerated as CFU/mL. **(C)** Differentiated HL-60 neutrophil-like cells were incubated with the strains shown at an MOI of 30, and intracellular bacterial loads determined after 24h. All experiments were performed in biological triplicate. Error bars represent ±SEM. Student’s t test with Welch’s correction was used to determine statistical significance relative to wildtype. ** P≤0.01,,***, P≤0.001.

### SSR42 is upregulated upon exposure to neutrophils via PerR derepression

It has previously been shown that *lukAB* expression is induced following exposure of *S. aureus* to neutrophils, and that this is mediated, indirectly, via the peroxide regulator PerR, which is known to derepress its targets in the presence of reactive oxygen species (31–33). Given that SSR42 plays an intimate role in intracellular survival and cytotoxicity, we next set out to determine if SSR42 was the missing link in this chain of regulation. To this end, we created a luciferase transcriptional reporter for the SSR42 promoter and exposed it to differentiated neutrophil-like HL-60 cells. In so doing, we determined that P_SSR42_ exhibited a >3-fold increase in activity after just 1h in the presence of neutrophils **(Fig. 3A)**. Next, to investigate if this regulation was governed by PerR, we re-assessed SSR42 promoter activity in a *perR* mutant. This time we determined that P_SSR42_ expression is significantly higher in the mutant strain when exposed to neutrophils as compared to wild-type **(Fig. 3B)**, which is consistent with PerR derepression.

**Figure 3.**
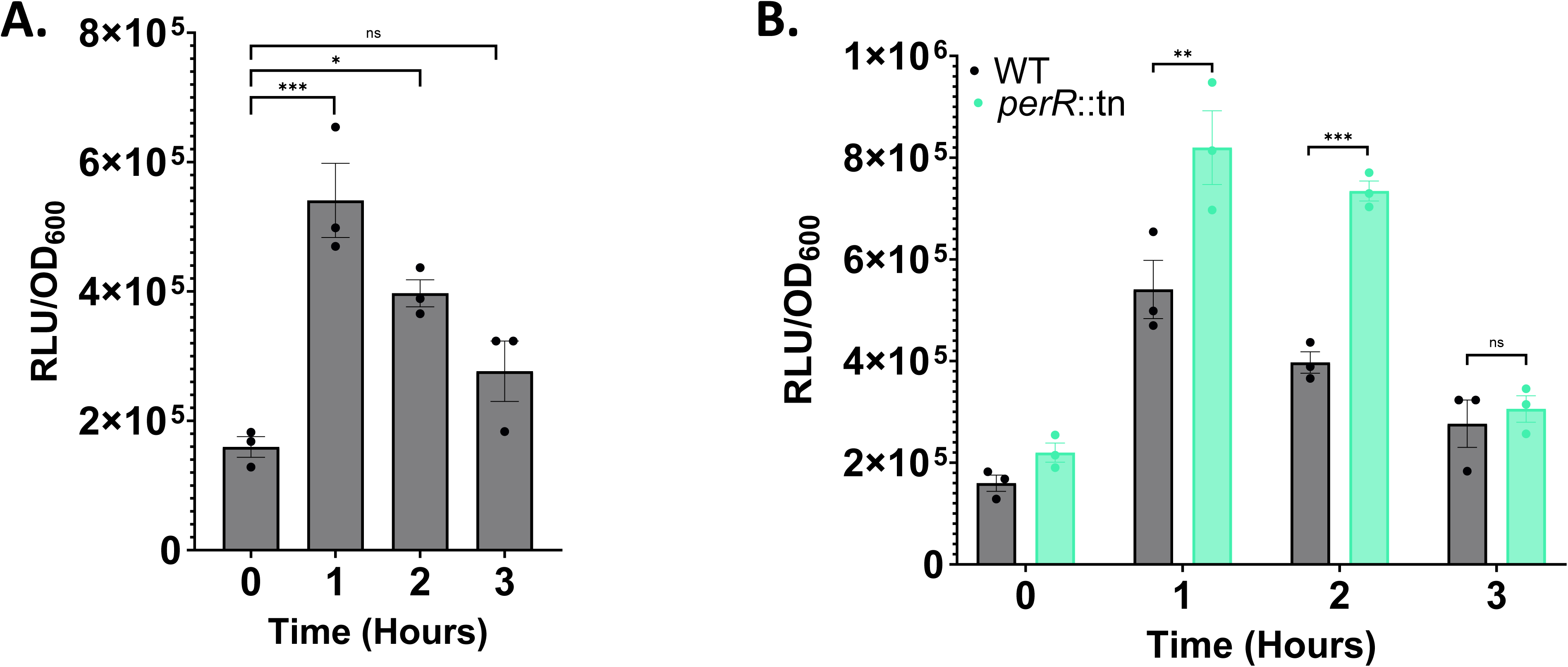
SSR42 is upregulated in response to neutrophils via PerR Derepression. **(A)** Differentiated HL60 cells were seeded at a concentration of 2×10^5^ before the *S. aureus* WT harboring a P_SSR42_-lux reporter fusion was added at a final OD_600_ of 0.1. Luminescence and OD_600_ reads were taken at T=0 and then hourly for 3 hours. Luminescence was normalized to OD_600_. Data is from biological triplicate samples. **(B)** As in A but comparing P_SSR42_ expression between the wild-type and a *perR* mutant strain. A student’s t-test with Welch’s correction was used to determine significance. Error bars are ±SEM. * P≤0.05 ** P≤0.01,,***, P≤0.001.

### SSR42 controls *lukAB* expression in response to neutrophils

We next set out to investigate whether SSR42 might mediate the impact of PerR on *lukAB* expression during engagement with neutrophils. As such, we created a P*_lukAB_-lux* fusion and assessed its response to neutrophils in the context of SSR42 and PerR. In so doing, we noted that upon addition of neutrophils, there was a significant increase in *lukAB* expression in the wild-type strain that mirrored that of P_SSR42_ **(Fig. 4A).** However, this response is ablated in ΔSSR42, where the luciferase signal is 5-fold lower in comparison to the wild-type after 2h of neutrophil exposure **(Fig. 4B).** To uncouple the effect of SSR42 from PerR, we next assessed *lukAB* expression in a *perR* mutant as well as an SSR42/*perR* double mutant. This revealed that, once again in a similar manner to SSR42, *lukAB* expression was higher in a *perR* mutant, however, *lukAB* expression in the SSR42/*perR* mutant mirrors that of ΔSSR42 **(Fig. 4 A-C)**. Native RNA levels were also quantified under these conditions, and identical induction patterns were observed **(Fig. S5**). This suggests that the response of *lukAB* to the presence of neutrophils is likely a result of direct regulation by SSR42, which is in turn controlled by PerR. This is further demonstrated upon overexpression of PerR in an SSR42 mutant, where there is no significant difference in P*_lukAB_* activity **(Fig. S6.)**.

**Figure 4.**
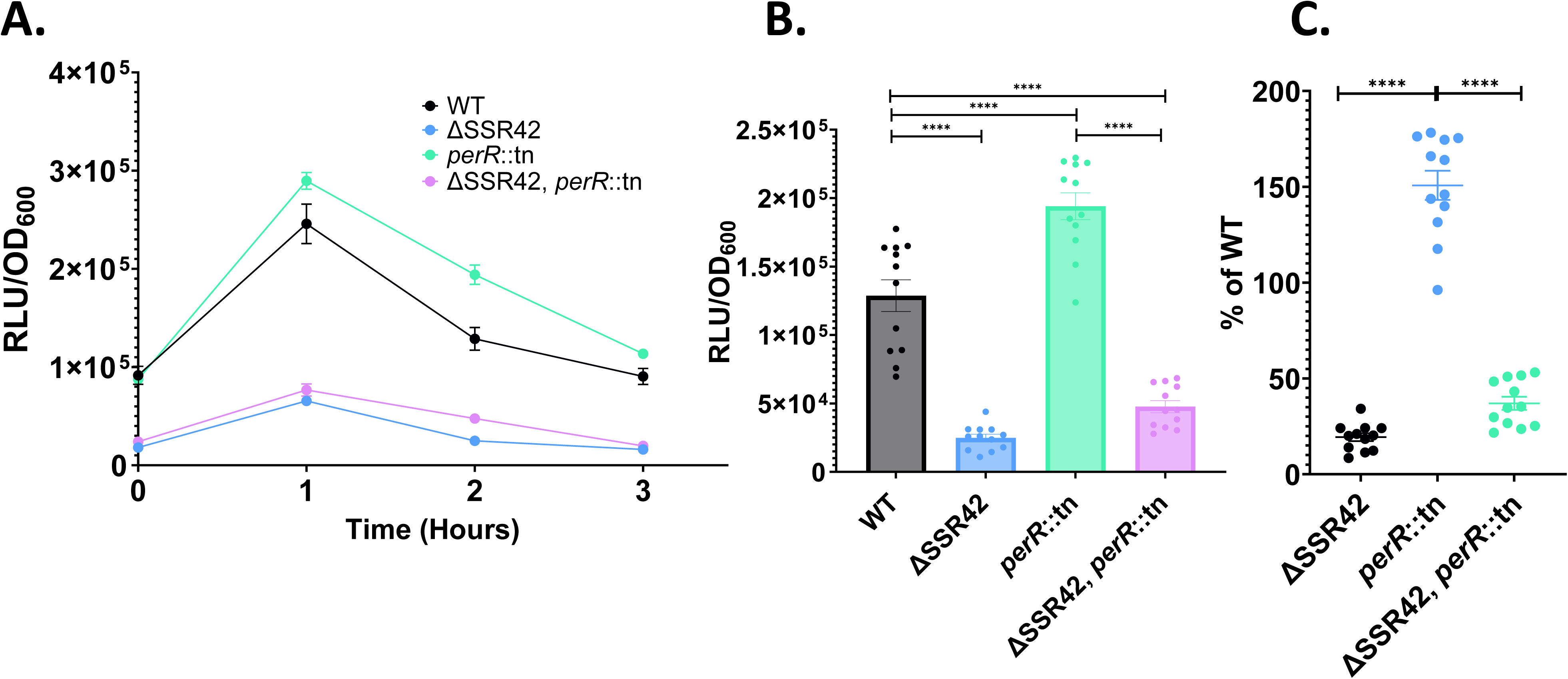
SSR42 controls LukAB expression in response to neutrophils. **(A)** As in Figure 3A but using the strains shown, all harboring a P*_lukAB_*-lux reporter fusion. **(B)** As in A but after 2h of exposure to neutrophils. **(C)** Data from B represented as a percentage of wild-type expression. Experiments were performed with 12 biological replicates. A one-way ANOVA was used to determine statistical significance. Error bars are ±SEM. ****, P<0.0001

### SSR42 regulates *lukAB* post-transcriptionally

To determine if the changes in *lukAB* expression are mediated by SSR42 binding to, and affecting stability of, the *lukAB* mRNA, we fused a fragment of the *lukA* 5’ UTR and a portion of its coding sequence in frame to GFP and expressed it under the control of a constitutive promoter in *S. aureus.* We then quantified GFP fluorescence in ΔSSR42 vs. wild-type, revealing a significant decrease in the stability of the translated LukA-GFP product in the mutant strain **(Fig. 5A).** Next, we used a two-plasmid system in which we expressed SSR42 under its native promoter alongside the LukA-GFP reporter in ΔSSR42. Here we observed a striking increase in LukA-GFP translation in the presence of SSR42 when compared to empty vector controls **(Fig. 5B).** Taken together, this suggests that SSR42 may bind post-transcriptionally to the *lukAB* mRNA to regulate its stability and thus translation.

**Figure 5.**
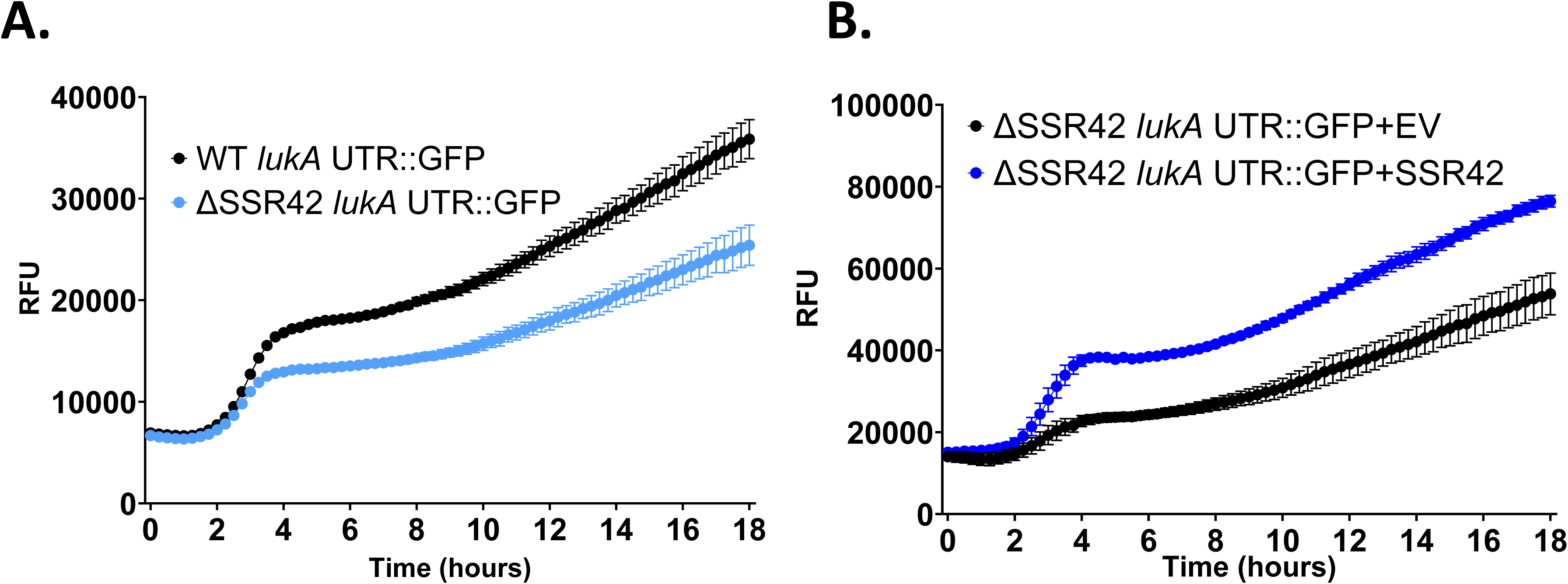
SSR42 post-transcriptionally controls LukAB abundance. **(A)** A portion of the *lukA* 5’ UTR and coding sequence was translationally fused to GFP, cloned into a shuttle vector using a constitutive promoter, and introduced into the wild-type or ΔSSR42 strains. **(B)** The *lukA-GFP* reporter fusion plasmid was co-expressed with either an empty vector (EV) or a plasmid containing SSR42 controlled by its native promoter in ΔSSR42. Fluorescence was measured every 15 minutes. Error bars are ± SEM. All experiments were performed in biological triplicate.

### The 3’ end of SSR42 is required for interaction with the *lukAB* 5’ UTR

To explore interaction between SSR42 and the *lukAB* mRNA further, we first used IntaRNA to predict possible base pairing interactions between the two RNA molecules (34). This predicted an interaction with the 3’ end of SSR42, ∼153bp from the end of the transcript, and the 5’ UTR of *lukAB*, ending 7bp upstream of the ribosome binding site (RBS) **(Fig. 6A)**. To investigate the relevance of this interaction *in vivo,* we first generated a series of truncated SSR42 variants and introduced these into ΔSSR42, before reassessing LukAB translation using the dual plasmid expression system. In so doing, we noted that shorter variants of SSR42 excluding the last 150-bp were unable to elicit wild-type levels of LukAB translation **(Fig. 6B).** This suggests that consistent with *in silico* predictions, the far 3’ end of SSR42 is primarily responsible for mediating interaction with *lukAB.* To next validate the binding region identified in *lukAB,* we performed mutagenesis on the predicted binding site within our translational LukAB-GFP fusion and assessed the impact of this on fluorescence **(Fig. 6C).** Upon disruption of the predicted binding sequence, we identify no difference between GFP levels in the wild-type and ΔSSR42, indicating that this region is indeed important for interaction with SSR42.

**Figure 6.**
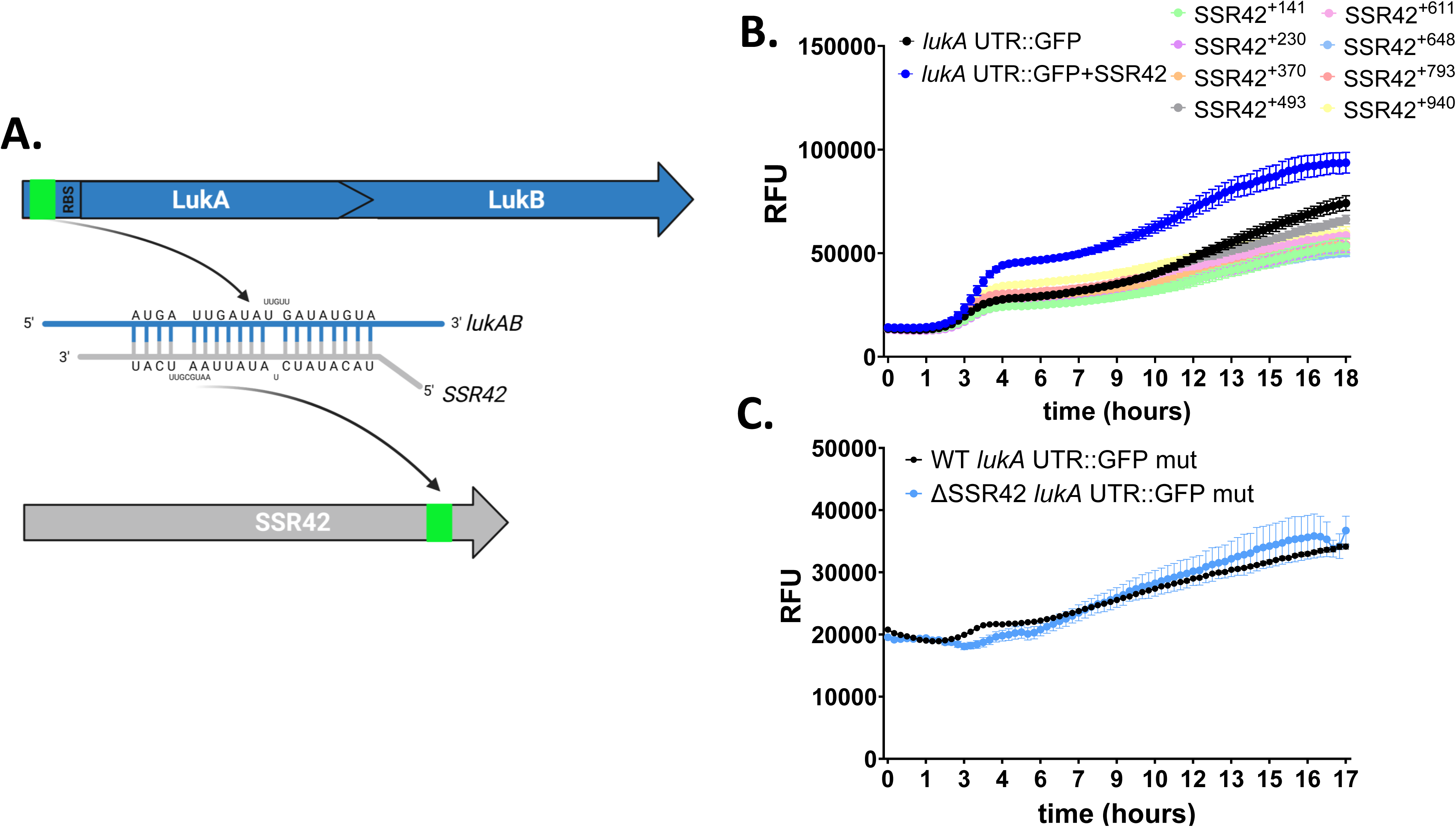
The 3’ end of SSR42 mediates binding to the *lukAB* 5’ UTR. **(A)** IntaRNA was used to identify thermodynamically likely regions of base paring between SSR42 and the *lukAB* mRNA. This identified an interaction region between the 3’ end of SSR42 and the 5’ UTR of *lukAB.* Created using Biorender.com. **(B)** The *lukA* UTR::GFP strain from Figure 5 was co-expressed in ΔSSR42 with either an empty vector (EV) or plasmids containing various truncated variants of SSR42. **(C)** The *lukA* UTR::GFP construct was mutagenized at the predicted site of interaction with SSR42 and luminescence was measured every 15 minutes in the wild-type and ΔSSR42. Error bars are ±SEM.

### SSR42 directly binds to the *lukAB* mRNA and enhances its stability

To explore further the direct interaction between SSR42 and the *lukAB* mRNA, electrophoretic mobility shift assays (EMSAs) were performed. A portion of the *lukA* 5’ UTR that included the region identified by IntaRNA was radiolabeled using *in vitro* transcription and incubated with increasing concentrations of *in vitro* transcribed SSR42. When these samples were co-incubated, we observed a concentration-dependent interaction between the two RNA molecules (**Fig. 7A).** By comparison, no binding was seen with radiolabeled *splE* mRNA under identical conditions, supporting the specificity of the SSR42-*lukAB* interaction. We next set out to investigate whether the observed binding had any impact on transcript stability. To this end, transcriptional arrest studies were performed on the wild type, mutant, and complemented strains grown for 15h. Samples were taken at t=0 before the addition of Rifampin, and then again at 5-, 10-, 30-, and 45-minutes posttreatment. RNA abundance was then quantified via qPCR, revealing that the *lukA* transcript degraded far more quickly, and to a greater overall degree, in the mutant strain when compared to the wild type and complement **(Fig. 7B).** When the mRNA half-life was calculated using a one phase decay curve, we found that the *lukAB* mRNA t_1/2_ was 2.9 minutes in ΔSSR42, but 4.5 minutes in the wild type. This suggests SSR42 is required for maintaining stability of the *lukAB* mRNA and facilitating translation of its protein products.

**Figure 7.**
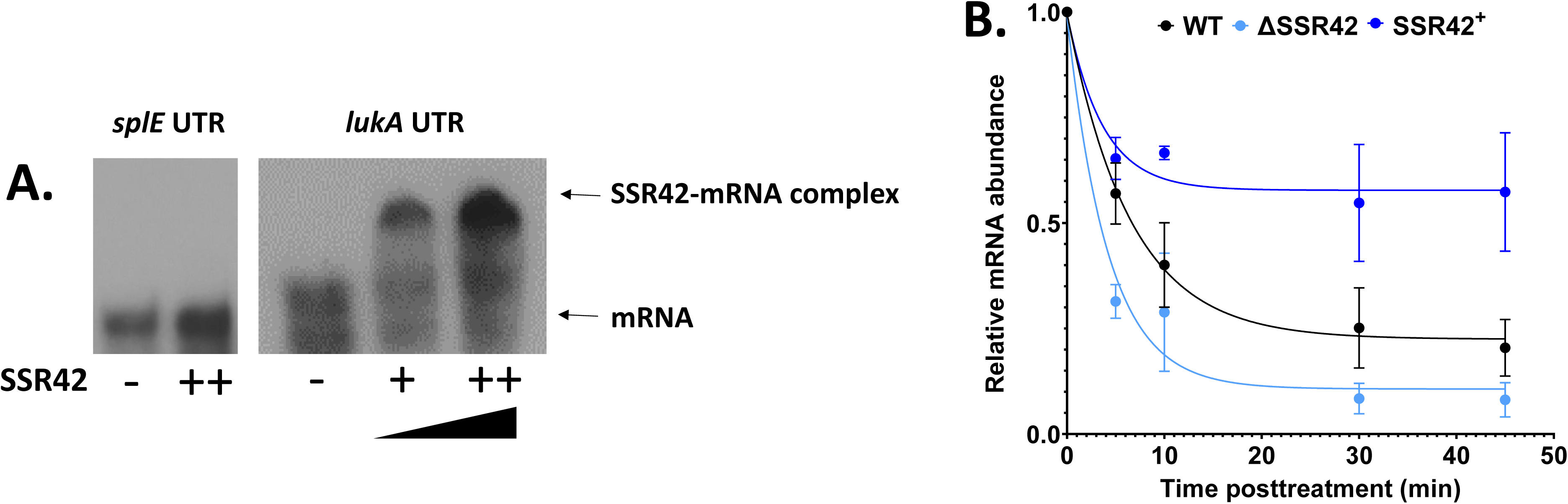
SSR42 directly binds and stabilizes the *lukAB mRNA*. **(A)** *In vitro* transcribed SSR42 was added in increasing amounts (1,000-fold to 1,500-fold excess) to radiolabeled portions of the *lukAB* mRNA. Samples were heated to 90° C for 2 min and then incubated at room temperature for 30 min before being loaded on to a non-denaturing gel. At identical concentrations, no binding is seen for radiolabeled *splE*. Images shown are representatives of 5 experimental replicates. **(B)** Bacterial cultures were grown in biological triplicate for 15 h before transcription was halted by the addition of 250 µg/mL Rifampin. Samples were collected 0, 5, 10, 30, and 45 minutes post-transcriptional arrest and *lukA* mRNA abundance was assessed via qPCR analysis. Data was compared to initial RNA abundance at t=0 and normalized to 16S rRNA. Shown is a one phase decay curve. Error bars are ±SEM.

### SSR42 is required to cause invasive *S. aureus* infections

Previous work has shown that SSR42 is important for localized infections using a murine model of abscess formation (26). To assess the role SSR42 plays in invasive disease we assessed the pathogenic capacity of the mutant in a murine model of septic infection. Mice were infected with ΔSSR42 or wild-type and the bacterial burden was determined across multiple organ systems after 7 days. Here we found that the mutant exhibited an abrogated ability to disseminate to the liver, heart, and spleen **(Fig. 8A).** Specifically, in the heart, ΔSSR42 exhibits a 4-log decrease in bacterial burden compared to WT, with 1-2-log decreases found in the liver and spleen. To explore this virulence defect more fully, we next used an acute model of murine pneumonia, intranasally inoculating mice with the wild type and ΔSSR42. After 24h of infection we assessed bacterial recovery from the lungs and once again found a major defect in the mutant’s ability to survive, with ΔSSR42 exhibiting a 3-log decrease in CFU/mL in comparison to the wild-type **(Fig. 8B)**. These findings, alongside those from others demonstrate that SSR42 is required for *S. aureus* infection in three different animal models, representing both localized and invasive forms of disease.

**Figure 8.**
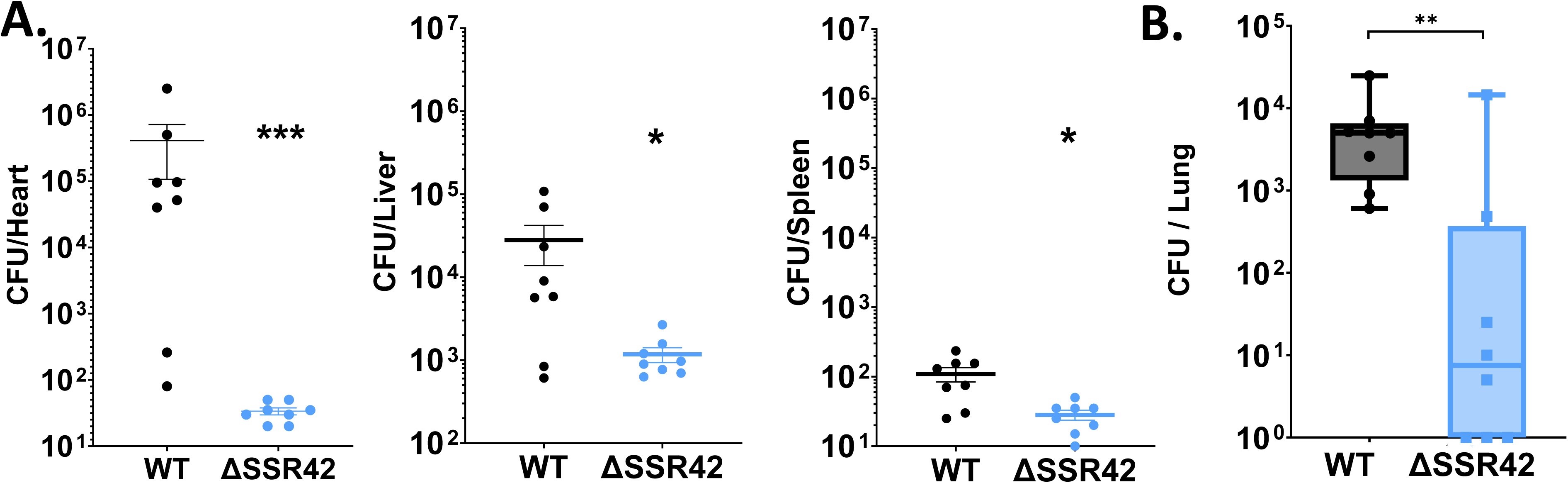
SSR42 is required for full virulence in murine models of invasive disease. **(A)** CD-1 mice were inoculated with the wild-type or ΔSSR42 at 5 x 10^8^ CFU via tail vein injection. Following 7 days of infection, organs were harvested and CFU determined. **(B)** C57BL/6J mice were intranasally inoculated with the wild-type or ΔSSR42 at 1 x 10^8^ CFU. After 24h, lungs were harvested and CFU determined. Whiskers represent minimum and maximum values. Solid lines represent median values. Mann Whitney tests were used to determine statistical significance relative to wildtype. * P≤0.05, **P≤0.01, ***P≤0.001. n=8 for each strain, error bars are ±SEM.

## Discussion

Recent years have seen an abundance of new discoveries regarding the complex nature of RNA-based regulation, revealing that many of these elements exhibit complex and nuanced roles in pathogenesis. Herein, we explore just such a factor, broadening the known regulome of the long noncoding RNA SSR42 from *S. aureus*. Using both transcriptomic and proteomic approaches we expand the SSR42 network of control almost three-fold from previous studies. From this, it is clear that the SSR42 regulome is far more complex than was previously ascertained. Of note, we identified no significant changes in any component relating to the SaeRS TCS, which previous studies suggested was responsible for SSR42-related regulation of at least some virulence factors (27). Instead, we note the strong downregulation of two SarA-homologs, *sarZ* and *sarR*, at - 9.6-fold and -62.6-fold, respectively, in ΔSSR42 compared to the wild-type. Both transcription factors have been characterized to play a role in promoting virulence factor expression in various *S. aureus* strains (35–38). Specifically, SarZ upregulates the expression of *aur, sspAB,* and *hla* while SarR activates *sspAB* and the *spl* proteases. Upon assessment of our own transcriptomics, the regulation of the proteases *aur* (via SarZ) and *sspAB* (via SarR) are consistent with the trends seen in our data, however the changes in *hla* (via SarZ) and *spl* (via SarR) expression are not. It is intriguing that these transcripts are so strikingly decreased in abundance in ΔSSR42, suggesting that SSR42 plays a key role in controlling their expression or stability. Indeed, bioinformatic searches for RNA-RNA binding do predict very strong interactions between SSR42 and the mRNAs of both transcription factors **(Fig. S7)**. Thus, it is possible that changes in *sarR* and *sarZ* expression may be causative for some, but not all, of the alterations in virulence factor transcription seen in the absence of SSR42.

When homing in on SSR42’s impact on the leukocidin LukAB, we show that SSR42 binding to the *lukAB* mRNA affects its stability, producing a concomitant impact on protein production. The latter effect, we posit, is reflective of this interaction altering the secondary structure of the *lukAB* mRNA to facilitate translation. Indeed, when predicting secondary structure for the *lukAB* mRNA we find that the RBS is predicted to lie locked within a hairpin, and thus may be inaccessible to the ribosome **(Fig. S8).** Just upstream of this lies the region where SSR42 appears to bind. Therefore, it is entirely plausible that binding of SSR42 5’ to the *lukA* RBS could alter the secondary structure such that the RBS is revealed and becomes accessible for translation. Of note, a very similar binding mechanism is seen for the well-studied interaction between RNAIII and the *hla* mRNA, whereby the former binds directly upstream of the latter’s RBS, liberating this sequence and activating translation (17). In the case of the *lukAB* mRNA, SSR42 binding also results in enhanced transcript stability, as in its absence we not only see decreased LukAB translation but also a significant decrease in *lukAB* mRNA abundance and half-life as well. Interestingly, in addition to the impact of SSR42 on *lukAB* mRNA stability and/or translational abundance, we also observe alterations in P*_lukAB_* promoter activity. This, at first glance, is contradictory to typical sRNA behavior, which usually only manifest post-transcriptionally. However, the P*_lukAB_* fragment used in our luciferase studies also contains the 5’ UTR region of the transcript, including the SSR42 binding site. Thus, as the absence of SSR42 destabilizes the *lukAB* mRNA, the promoter fusion transcript will inevitably be destabilized in our luciferase assays as well.

Recent work by Savin et al. identified the peroxide sensor PerR as a regulator of LukAB activity that responds to the presence of hPMNs and leads to enhanced leukocidin expression (33). Their study suggested no direct regulation between PerR and *lukAB* as there are no PerR binding boxes detected in the leukocidin promoter. In our study, we observed that the SSR42 promoter is also induced in the presence of human neutrophil-like cells and, importantly, is required for the *lukAB* promoter to respond under these conditions. Investigation of P_SSR42_ architecture revealed 3 putative PerR binding boxes (sequence identified by Horsburgh et. al (32)) upstream and/or within the SSR42 transcript **(Fig. S9)**. It has previously been shown that known PerR targets such as *katA, ahpC,* and *ftnA* can possess multiple binding boxes within their promoters and are de-repressed during oxidative stress (31–33). When we mutated each of these binding sequences individually, we identified no changes in promoter induction (data not shown), however, we were unable to combinatorially mutate more than one box within the promoter sequence at the same time. It is possible that profound derepression in the absence of these PerR binding sites results in toxic over production of SSR42. Beyond this, there are likely multiple factors controlling the expression of SSR42 as its production is still modulated in response to neutrophils in the absence of PerR. We should note that these findings were observed using differentiated HL-60 cells rather than primary human neutrophils, thus, additional studies using primary *ex vivo* samples would be highly valuable for corroborating our findings. Moreover, to further test for the full nuance of SSR42 induction within the host, it would be intriguing to investigate how inhibition of the oxidative burst of neutrophils within murine systems impacts SSR42 production.

The promoter of SSR42, as well as its primary regulator Rsp, have previously been shown to be induced in response to the presence of hydrogen peroxide (39). Moreover, previous work identified that Rsp is required for full cytotoxicity towards epithelial cells and neutrophils in an Agr-independent manner (39). We posit that these phenotypes are at least partially SSR42-mediated. To explore this, we assessed expression from the *rsp* promoter in the presence of neutrophils and found that it too is induced by their presence **(Fig. S10)**. Importantly, however, P*_rsp_* is not derepressed by PerR in this context, suggesting that the PerR-mediated control of *lukAB* by SSR42 is independent of Rsp. This demonstrates that, depending on the environment, Rsp is necessary, but not sufficient, for SSR42 expression. Thus, we propose that SSR42 is the missing link between PerR-mediated ROS sensing and subsequent activation of LukAB activity to mediate survival and escape from phagocytic cells. Further evidence for PerR-mediated regulation of SSR42 also arises from a recent study by our group. Here, upon assessing transcriptomic changes in response to treatment with Calprotectin (i.e. metal starved), SSR42 was found to be downregulated ∼2-fold. When samples were treated with a mutated form of Calprotectin that only sequesters Zn and Cu, SSR42 was upregulated >3-fold (40). In *S. aureus,* PerR requires Fe and Mn for oxidation in response to endogenous H_2_O_2_ (31); thus, it follows that derepression of SSR42 would be detected only in conditions including these metals. Finally, it is intriguing to consider that P_SSR42_ may also be activated in response to bacterial-intrinsic ROS, as SSR42 is highly abundant in stationary phase without the presence of host cells. Indeed, PerR derepression has been shown to occur in response to intrinsic H_2_O_2_ levels, which are highest in stationary phase *S. aureus* cells (32).

Metal homeostasis and the PerR-mediated oxidative stress response are intimately linked through a network of metalloregulators: Fur, Zur, and MntR (32, 41–44). Interestingly, the *fur* transcript is upregulated 3.14-fold in our SSR42 mutant while *mntR* expression is decreased -2-fold. It is important to note that in addition to regulating SSR42, PerR plays a role in regulating Fur and MntR expression, while all three players act in a coordinated manner to induce overlapping regulons contributing to oxidative stress resistance, including factors such as catalase (KatA), thioredoxin reductase (TrxB), and alkylhydroperoxide reductase (AhpCF) (41, 44). While these elements were not altered in the SSR42 mutant transcriptome, we did uncover three alternative antioxidant-like factors that were decreased in expression in the SSR42 mutant. These include a nitroreductase linked to iron acquisition (*ntrA,* -3.41-fold), a nitroreductase with glutathione antioxidant behavior (SAUSA300_0790, -2.26-fold), and a thioredoxin-like protein (SAUSA300_0795, -2.27-fold) (45–47). Thus, we posit that SSR42 weaves a common thread connecting these metallo-oxidative regulators and subsequently impacts virulence factor abundance in response to ROS-generating niches. Key examples of this behavior in *S. aureus* have been shown in the case of IsrR and RsaC, two sRNAs that are expressed in low iron or manganese environments, respectively, and function to enhance the oxidative stress response and pathogenesis upon activation (25, 48–50).

When considering the wider context of *lukAB* regulation, it is strikingly diverse, and not limited merely to induction in the presence of neutrophils and/or reactive oxygen species. For example, as with numerous other exotoxins, alterations in LukAB abundance have been linked to the Agr system through toxin suppression via Rot; although there is currently no evidence to show any direct interaction between Rot and P*_lukAB_* (51). SaeRS, which has been shown to be crucial for pathogenesis and survival within neutrophils, serves as a primary regulator of LukAB abundance through proposed binding at the promoter region (52–54). Additionally, the ArlRS system has been shown to impact LukAB abundance likely through its effector MgrA; although once again no direct interactions have been identified (55). Finally, the autolysin Atl indirectly controls LukAB trafficking and release through modulation of cell wall architecture via cleavage of peptidoglycan (56). Collectively, despite the fact that LukAB has been shown to be vital for pathogenesis and interaction with components of host immunity, the full regulatory network that controls its expression has yet to be fully elucidated.

Taken together, our findings expand the role of SSR42 beyond a direct virulence-related pathway and illustrate a much more complex role for this sRNA as it pertains to sensing oxidative stress, nutritional immunity, and other exogenous signals. A proposed mechanism of SSR42’s involvement in the regulation of these processes as well as its role in the broader regulation of LukAB is summarized in **Figure 9**. To our knowledge, this is the first direct RNA-based regulation of *lukAB* characterized to date in *S. aureus*. Not only do we uncover a novel regulator or leukocidin activity, but we begin to unravel the labyrinthine network that controls the response to changing environments, particularly within the host. Indeed, past work on SSR42 demonstrated its role in murine abscess formation, while our own study broadens this to include complex and disseminated infections including sepsis and pneumonia. Thus, we present SSR42 as a global regulatory RNA that responds to pathophysiologically relevant conditions to bolster virulence factor abundance and facilitate pathogenesis in *S. aureus*.

**Figure 9.**
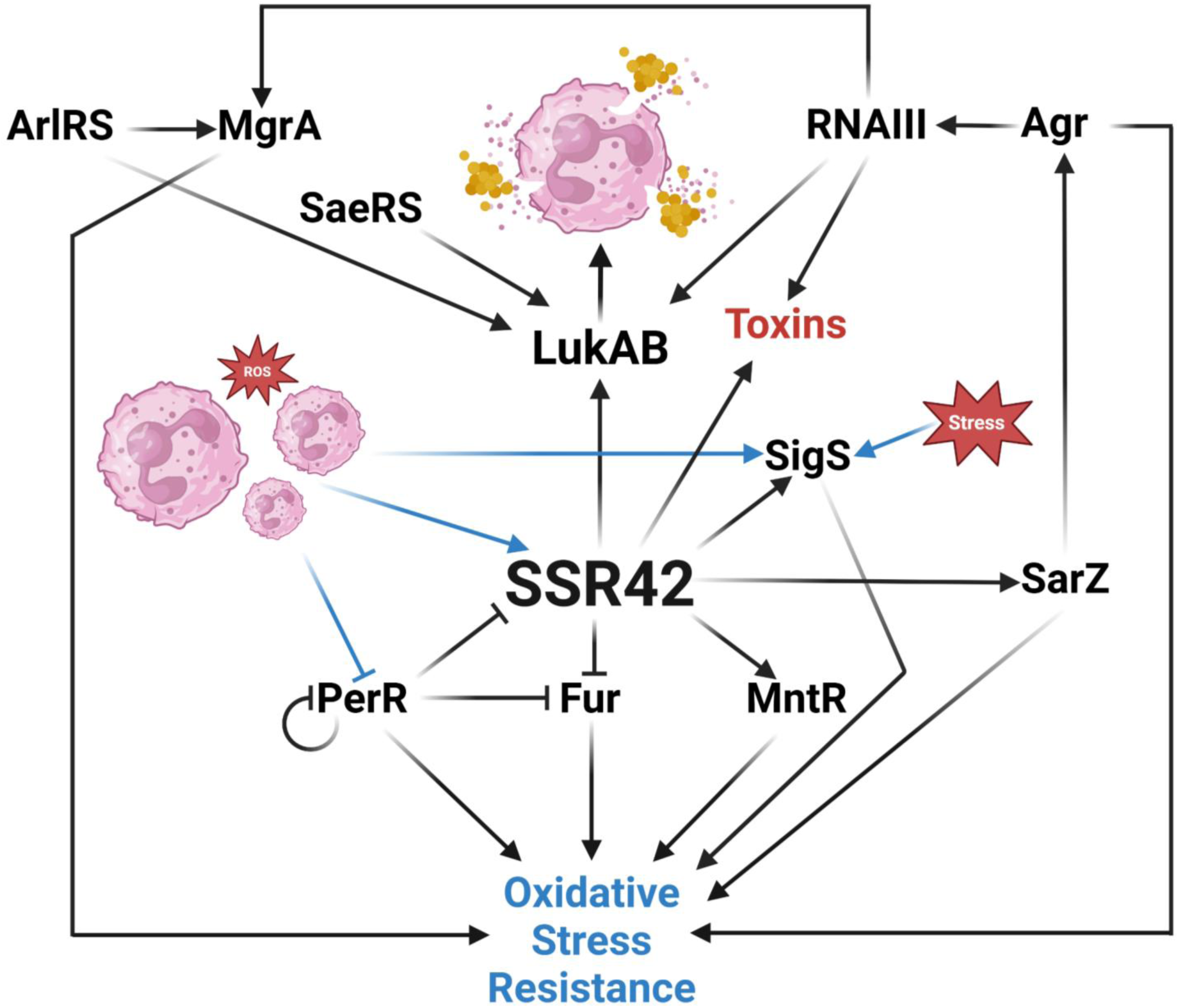
Proposed role of SSR42 in *S. aureus*. The presence of neutrophils and/or reactive oxygen species leads to PerR-mediated derepression of SSR42, which then binds and stabilizes *lukAB* mRNA in order to defend against phagocytosis and/or facilitate escape. The regulatory links between the metalloregulators, stress response, and oxidative stress resistance is represented alongside known regulation of the LukAB and virulence. Created with BioRender.com

## Supporting information

Combined supplementary methods, tables, and figures

## Acknowledgements

This study was supported by grants AI124458 and AI157506 (both to L.N.S.) from the National Institute of Allergy and Infectious Diseases.

## References

1. Carroll RK, Weiss A, Broach WH, Wiemels RE, Mogen AB, Rice KC, Shaw LN. 2016. Genome-wide annotation, identification, and global transcriptomic analysis of regulatory or small RNA gene expression in *Staphylococcus aureus*. mBio 7.

2. Liu W, Rochat T, Toffano-Nioche C, Le Lam TN, Bouloc P, Morvan C. 2018. Assessment of bona fide sRNAs in *Staphylococcus aureus*. Front Microbiol 9:1– 13.

3. Pichon C, Felden B. 2005. Small RNA genes expressed from *Staphylococcus aureus* genomic and pathogenicity islands with specific expression among pathogenic strains. Proc Natl Acad Sci U S A 102:14249–14254.

4. Laner EA, Carroll K. 2020. Reading between the Lines: Utilizing RNA-Seq Data for Global Analysis of sRNAs in *Staphylococcus aureus*. mSphere 5(4):e00439–20.

5. Broach WH, Weiss A, Shaw LN. 2016. Transcriptomic analysis of staphylococcal sRNAs: Insights into species-specific adaption and the evolution of pathogenesis. Microb Genomics 2 (7).

6. Bronsard J, Pascreau G, Sassi M, Mauro T, Augagneur Y, Felden B. 2017. SRNA and cis-Antisense sRNA identification in *Staphylococcus aureus* highlights an unusual sRNA gene cluster with one encoding a secreted peptide. Sci Rep 7:1– 17.

7. Menard G, Silard C, Suriray M, Rouillon A, Augagneur Y. 2022. Thirty Years of sRNA-Mediated Regulation in *Staphylococcus aureus*: From Initial Discoveries to In Vivo Biological Implications. International Journal of Molecular Sciences. 23(13), 7346.

8. Barrientos L, Mercier N, Lalaouna D, Caldelari I. 2021. Assembling the Current Pieces: The Puzzle of RNA-Mediated Regulation in *Staphylococcus aureus*. Front Microbiol 12:1–7.

9. Bastock RA, Marino EC, Wiemels RE, Holzschu DL, Keogh RA, Zapf RL, Murphy ER, Carroll RK. 2021. *Staphylococcus aureus* Responds to Physiologically Relevant Temperature Changes by Altering Its Global Transcript and Protein Profile. mSphere 6.

10. Bordeau V, Cady A, Revest M, Rostan O, Sassi M, Tattevin P, Donnio PY, Felden B. 2016. *Staphylococcus aureus* regulatory RNAs as potential biomarkers for bloodstream infections. Emerg Infect Dis 22:1570–1578.

11. Kim S, Reyes D, Beaume M, Francois P, Cheung A. 2014. Contribution of teg49 Small RNA in the 5’ Upstream Transcriptional Region of sarA to Virulence in *Staphylococcus aureus*. Infect Immun 82:4369–4379.

12. Le Huyen KB, Gonzalez CD, Pascreau G, Bordeau V, Cattoir V, Liu W, Bouloc P, Felden B, Chabelskaya S. 2021. A small regulatory RNA alters *Staphylococcus aureus* virulence by titrating RNAIII activity. Nucleic Acids Res 49:10644–10656.

13. McKellar SW, Ivanova I, Arede P, Zapf RL, Mercier N, Chu LC, Mediati DG, Pickering AC, Briaud P, Foster RG, Kudla G, Ross Fitzgerald J, Caldelari I, Carroll RK, Tree JJ, Granneman S. 2022. RNase III CLASH in MRSA uncovers sRNA regulatory networks coupling metabolism to toxin expression. Nat Commun 13.

14. Gupta RK, Luong TT, Lee CY. 2015. RNAIII of the *Staphylococcus aureus* agr system activates global regulator MgrA by stabilizing mRNA. Proc Natl Acad Sci U S A 112:14036–14041.

15. Huntzinger E, Boisset S, Saveanu C, Benito Y, Geissmann T, Namane A, Lina G, Etienne J, Ehresmann B, Ehresmann C, Jacquier A, Vandenesch F, Romby P. 2005. *Staphylococcus aureus* RNAIII and the endoribonuclease III coordinately regulate spa gene expression. EMBO Journal 24:824–835.

16. Boisset S, Geissmann T, Huntzinger E, Fechter P, Bendridi N, Possedko M, Chevalier C, Helfer AC, Benito Y, Jacquier A, Gaspin C, Vandenesch F, Romby P. 2007. *Staphylococcus aureus* RNAIII coordinately represses the synthesis of virulence factors and the transcription regulator Rot by an antisense mechanism. Genes Dev 21:1353–1366.

17. Morfeldt E, Taylor D, Von Gabain A, Arvidson S. 1995. Activation of alpha-toxin translation in *Staphylococcus aureus* by the trans-encoded antisense RNA, RNAIII. EMBO Journal 14:4569–4577.

18. Briaud P, Zapf RL, Mayher AD, McReynolds AKG, Frey A, Sudnick EG, Wiemels RE, Keogh RA, Shaw LN, Bose JL, Carroll RK. 2022. The Small RNA Teg41 Is a Pleiotropic Regulator of Virulence in *Staphylococcus aureus*. Infect Immun 19.

19. Zapf RL, Wiemels RE, Keogh RA, Holzschu DL, Howell KM, Trzeciak E, Caillet AR, King KA, Selhorst SA, Naldrett MJ, Bose JL, Carroll RK. 2019. The small RNA Teg41 regulates expression of the alpha phenol-soluble modulins and is required for virulence *in Staphylococcus aureus*. mBio 10:1–19.

20. Manna AC, Leo S, Girel S, González-Ruiz V, Rudaz S, Francois P, Cheung AL. 2022. Teg58, a small regulatory RNA, is involved in regulating arginine biosynthesis and biofilm formation in *Staphylococcus aureus*. Sci Rep 12:1–18.

21. Zhou J, Zhao H, Yang H, He C, Shu W, Cui Z, Liu Q. 2022. Insights Into the Impact of Small RNA SprC on the Metabolism and Virulence of *Staphylococcus aureus*. Front Cell Infect Microbiol 12:1–13.

22. Le Huyen KB, Gonzalez CD, Pascreau G, Bordeau V, Cattoir V, Liu W, Bouloc P, Felden B, Chabelskaya S. 2021. A small regulatory RNA alters *Staphylococcus aureus* virulence by titrating RNAIII activity. Nucleic Acids Res 49:10644–10656.

23. Chabelskaya S, Gaillot O, Felden B. 2010. A *Staphylococcus aureus* small RNA is required for bacterial virulence and regulates the expression of an immune-evasion molecule. PLoS Pathog 6:1–11.

24. Chabelskaya S, Bordeau V, Felden B. 2014. Dual RNA regulatory control of a *Staphylococcus aureus* virulence factor. Nucleic Acids Res 42:4847–4858.

25. Lalaouna D, Baude J, Wu Z, Tomasini A, Chicher J, Marzi S, Vandenesch F, Romby P, Caldelari I, Moreau K. 2019. RsaC sRNA modulates the oxidative stress response of *Staphylococcus aureus* during manganese starvation. Nucleic Acids Res 47:9871–9887.

26. Morrison JM, Miller EW, Benson MA, Alonzo F, Yoong P, Torres VJ, Hinrichs SH, Dunmana PM. 2012. Characterization of SSR42, a Novel virulence factor regulatory RNA that contributes to the pathogenesis of a *Staphylococcus aureus* USA300 representative. J Bacteriol 194:2924–2938.

27. Horn, J., Klepsch, M., Manger, M., Wolz, C., Rudel, T., & Fraunholz, M. (2018). Long Noncoding RNA SSR42 Controls *Staphylococcus aureus* Alpha-Toxin Transcription in Response to Environmental Stimuli. Journal of bacteriology, 200(22), e00252–18.

28. DuMont AL, Nygaard TK, Watkins RL, Smith A, Kozhaya L, Kreiswirth BN, Shopsin B, Unutmaz D, Voyich JM, Torres VJ. 2011. Characterization of a new cytotoxin that contributes to *Staphylococcus aureus* pathogenesis. Mol Microbiol 79:814–825.

29. DuMont AL, Yoong P, Day CJ, Alonzo F, McDonald WH, Jennings MP, Torres VJ. 2013. *Staphylococcus aureus* LukAB cytotoxin kills human neutrophils by targeting the CD11b subunit of the integrin Mac-1. Proc Natl Acad Sci U S A 110:10794–10799.

30. DuMont AL, Yoong P, Surewaard BGJ, Benson MA, Nijland R, Strijp JAG van, Torres VJ. 2013. *Staphylococcus aureus* elaborates leukocidin AB to mediate escape from within human neutrophils. Infect Immun 81:1830–1841.

31. Ji CJ, Kim JH, Won Y Bin, Lee YE, Choi TW, Ju SY, Youn H, Helmann JD, Lee JW. 2015. *Staphylococcus aureus* PerR is a hypersensitive hydrogen peroxide sensor using iron-mediated histidine oxidation. Journal of Biological Chemistry 290:20374–20386.

32. Horsburgh MJ, Clements MO, Crossley H, Ingham E, Foster SJ. 2001. PerR controls oxidative stress resistance and iron storage proteins and is required for virulence in *Staphylococcus aureus*. Infect Immun 69:3744–3754.

33. Savin, A., Anderson, E. E., Dyzenhaus, S., Podkowik, M., Shopsin, B., Pironti, A., & Torres, V. J. 2024. *Staphylococcus aureus* senses human neutrophils via PerR to coordinate the expression of the toxin LukAB. Infection and immunity, 92(2), e0052623.

34. Mann M, Wright PR, Backofen R. 2017. IntaRNA 2.0: Enhanced and customizable prediction of RNA-RNA interactions. Nucleic Acids Res 45:W435–W439.

35. Manna A, Cheung AL. 2001. Characterization of sarR, a modulator of sar expression in *Staphylococcus aureus*. Infect Immun 69:885–896.

36. Reyes D, Andrey DO, Monod A, Kelley WL, Zhang G, Cheung AL. 2011. Coordinated regulation by AgrA, SarA, and SarR to control agr expression in *Staphylococcus aureus*. J Bacteriol 193:6020–6031.

37. Tamber S, Cheung AL. 2009. SarZ promotes the expression of virulence factors and represses biofilm formation by modulating SarA and agr in *Staphylococcus aureus*. Infect Immun 77:419–428.

38. Chen PR, Nishida S, Poor CB, Cheng A, Bae T, Kuechenmeister L, Dunman PM, Missiakas D, He C. 2009. A new oxidative sensing and regulation pathway mediated by the MgrA homologue SarZ in *Staphylococcus aureus*. Mol Microbiol 71:198–211.

39. Das S, Lindemann C, Young BC, Muller J, Österreich B, Ternette N, Winkler AC, Paprotka K, Reinhardt R, Förstner KU, Allen E, Flaxman A, Yamaguchi Y, Rollier CS, Diemen P Van, Blättner S, Remmele CW, Selle M, Dittrich M, Müller T, Vogel J, Ohlsen K, Crook DW, Massey R, Wilson DJ, Rudel T, Wyllie DH, Fraunholz MJ. 2016. Natural mutations in a *Staphylococcus aureus* virulence regulator attenuate cytotoxicity but permit bacteremia and abscess formation. Proc Natl Acad Sci U S A 113:E3101–E3110.

40. Ruiz VM, Freiberg JA, Weiss A, Green ER, Jobson M, Felton E, Shaw LN, Chazin WJ, Skaar EP. 2024. Coordinated adaptation of *Staphylococcus aureus* to calprotectin-dependent metal sequestration. mBio 15:e01389–24.

40. Horsburgh MJ, Ingham E, Foster SJ. 2001. In *Staphylococcus aureus*, Fur is an interactive regulator with PerR, contributes to virulence, and is necessary for oxidative stress resistance through positive regulation of catalase and iron homeostasis. J Bacteriol 183:468–475.

41. Torres VJ, Attia AS, Mason WJ, Hood MI, Corbin BD, Beasley FC, Anderson KL, Stauff DL, McDonald WH, Zimmerman LJ, Friedman DB, Heinrichs DE, Dunman PM, Skaar EP. 2010. *Staphylococcus aureus* fur regulates the expression of virulence factors that contribute to the pathogenesis of pneumonia. Infect Immun 78:1618–1628.

42. Lindsay JA, Foster SJ. 2001. Zur: A Zn2+-responsive regulatory element of *Staphylococcus aureus*. Microbiology (N Y) 147:1259–1266.

43. Horsburgh MJ, Wharton SJ, Cox AG, Ingham E, Peacock S, Foster SJ. 2002. MntR modulates expression of the PerR regulon and superoxide resistance in *Staphylococcus aureus* through control of manganese uptake. Mol Microbiol 44:1269–1286.

44. Tavares AFN, Nobre LS, Melo AMP, Saraiva LM. 2009. A novel nitroreductase of *Staphylococcus aureus* with S-nitrosoglutathione reductase activity. J Bacteriol 191:3403–3406.

45. Liochev SI, Hausladen A, Fridovich I. 1999. Nitroreductase A is regulated as a member of the soxRS regulon of *Escherichia coli*. Proc Natl Acad Sci U S A 96:3537–3539.

46. Nishinaka Y, Masutani H, Nakamura H, Yodoi J. 2001. Regulatory roles of thioredoxin in oxidative stress-induced cellular responses. Redox Report 6:289– 295.

47. Ganske A, Busch LM, Hentschker C, Reder A, Michalik S, Surmann K, Völker U, Mäder U. 2024. Exploring the targetome of IsrR, an iron-regulated sRNA controlling the synthesis of iron-containing proteins in *Staphylococcus aureus* 1– 19.

48. Coronel-Tellez RH, Pospiech M, Barrault M, Liu W, Bordeau V, Vasnier C, Felden B, Sargueil B, Bouloc P. 2022. sRNA-controlled iron sparing response in Staphylococci. Nucleic Acids Res 50:8529–8546.

49. Rios-Delgado G, McReynolds AKG, Pagella EA, Norambuena J, Briaud P, Zheng V, Munneke MJ, Kim J, Racine H, Carroll R, Zelzion E, Skaar E, Bose JL, Parker D, Lalaouna D, Boyd JM. 2024. The *Staphylococcus aureus* small non-coding RNA IsrR regulates TCA cycle activity and virulence. bioRxiv 2024.07.03.601953.

50. Mootz JM, Benson MA, Heim CE, Crosby HA, Kavanaugh JS, Dunman PM, Kielian T, Torres VJ, Horswill AR. 2015. Rot is a key regulator of *Staphylococcus aureus* biofilm formation. Mol Microbiol 96:388–404.

51. Voyich, J. M., Vuong, C., DeWald, M., Nygaard, T. K., Kocianova, S., Griffith, S., Jones, J., Iverson, C., Sturdevant, D. E., Braughton, K. R., Whitney, A. R., Otto, M., & DeLeo, F. R. 2009. The SaeR/S gene regulatory system is essential for innate immune evasion by *Staphylococcus aureus*. The Journal of infectious diseases, 199(11), 1698–1706.

52. Nygaard, T. K., Pallister, K. B., Ruzevich, P., Griffith, S., Vuong, C., & Voyich, J. M. (2010). SaeR binds a consensus sequence within virulence gene promoters to advance USA300 pathogenesis. The Journal of infectious diseases, 201(2), 241– 254.

53. Flack CE, Zurek OW, Meishery DD, Pallister KB, Malone CL, Horswill AR, Voyich JM. 2014. Differential regulation of staphylococcal virulence by the sensor kinase SaeS in response to neutrophil-derived stimuli. Proc Natl Acad Sci U S A 111:2037–2045.

54. Crosby HA, Tiwari N, Kwiecinski JM, Xu Z, Dykstra A, Jenul C, Fuentes EJ, Horswill AR. 2020. The *Staphylococcus aureus* ArlRS two-component system regulates virulence factor expression through MgrA. Mol Microbiol 113:103–122.

55. Zheng X, Ma SX, John AS, Torres VJ. 2022. The Major Autolysin Atl Regulates the Virulence of *Staphylococcus aureus* by Controlling the Sorting of LukAB. Infect Immun 90:1–13.

