## Supplementary material for "SSR42 is a Novel Regulator of Cytolytic Activity in *Staphylococcus aureus*": Combined supplementary methods, tables, and figures

Running title: SSR42 controls *Staphylococcus aureus* cytotoxicity.

### Supplemental Methods

**Bacterial strains and growth conditions:** All bacterial strains and plasmids used for this study are listed in **Supplemental Table S2** and **Supplemental Table S3**, respectively. Overnight cultures were routinely grown at 37°C with shaking at 250 rpm in 5mL of tryptic soy broth (TSB) for *S. aureus* or lysogeny broth (LB) for *E. coli.* When required, media was supplemented with the following antibiotics: for *E. coli*: 100μg/mL ampicillin; for *S. aureus*: 10µg/mL chloramphenicol, 5µg/mL erythromycin, 25µg/mL lincomycin, and 5µg/mL tetracycline. To obtain synchronous growth of *S. aureus,* overnight cultures were diluted 1:100 into fresh TSB and grown for 3h before being standardized to a uniform $OD_{600}$ of 0.05 in fresh TSB. Unless otherwise described, cell pellets were harvested via centrifugation at 4000 x *g* for 10 minutes.

**Supplemental Table S2.** Bacterial strains used in this study.

| Strain Name | Description | Source |
|---|---|---|
| ***E. coli*** | | |
| DH5α | Cloning strain | (1) |
| JM109 | Cloning strain | Promega pGEM-T Easy Vector Kit |
| MEJ3834 | JM109 pGEM-T::*lukA* UTR | This study |
| MEJ3845 | JM109 pGEM-T::SSR42 | This study |
| ***S. aureus*** | | |
| RN4220 | Cloning strain, restriction-deficient | Lab stock |
| TCH1516 | USA300 HOU CA-MRSA $Erm^{s}$, Wildtype parent strain | (2) |
| BRT2544 | TCH1516 ΔSSR42 | This study |
| NE665 | JE2 *perR*::tn | (3) |

| | | |
|---|---|---|
| MEJ4002 | TCH1516 *perR*::tn | This study |
| MEJ4003 | TCH1516 ΔSSR42, *perR*::tn | This study |
| LNS1927 | TCH1516 pMK4 EV | This study |
| BRT2545 | TCH1516 ΔSSR42 pMK4 EV | This study |
| BRT2546 | TCH1516 ΔSSR42 pMK4::SSR42 | This study |
| MEJ4000 | TCH1516 pXen-1::$P_{SSR42}$ | This study |
| MEJ4001 | TCH1516 *perR*::tn, pXen-1::$P_{SSR42}$ | This study |
| MEJ3546 | TCH1516 pXen-1::$P_{lukAB}$ | This study |
| MEJ3550 | TCH1516 ΔSSR42 pXen-1::$P_{lukAB}$ | This study |
| MEJ3826 | TCH1516 *perR*::tn, pXen-1::$P_{lukAB}$ | This study |
| MEJ3827 | TCH1516 ΔSSR42, *perR*::tn, pXen-1::$P_{lukAB}$ | This study |
| MEJ3828 | TCH1516 ΔSSR42, *perR*::tn, pXen-1::$P_{lukAB}$, pJB67::PerR | This study |
| MEJ3831 | TCH1516 pCN33::*lukA* UTR | This study |
| MEJ3832 | TCH1516 ΔSSR42 pCN33::*lukA* UTR | This study |
| MEJ3837 | +SSR42, ΔSSR42 pICS3::SSR42 | This study |
| MEJ3838 | +SSR42$^{+141}$, ΔSSR42 pICS3::141-nt SSR42 | This study |
| MEJ3839 | +SSR42$^{+230}$, ΔSSR42 pICS3::230-nt SSR42 | This study |

| | | |
|---|---|---|
| MEJ3840 | +SSR42$^{+370}$, ΔSSR42 pICS3::370-nt SSR42 | This study |
| MEJ3841 | +SSR42$^{+493}$, ΔSSR42 pICS3::493-nt SSR42 | This study |
| MEJ3842 | +SSR42$^{+611}$, ΔSSR42 pICS3::611-nt SSR42 | This study |
| MEJ3843 | +SSR42$^{+648}$, ΔSSR42 pICS3::648-nt SSR42 | This study |
| MEJ3844 | +SSR42$^{+793}$, ΔSSR42 pICS3::793-nt SSR42 | This study |
| MEJ3845 | +SSR42$^{+940}$, ΔSSR42 pICS3::940-nt SSR42 | This study |
| MEJ4022 | TCH1516 pCN33::*lukA*::GFP MUT | This study |

**Supplemental Table S3.** Bacterial strains and plasmids used in this study.

| Plasmids | Description | Source |
|---|---|---|
| pJB38 | Allelic replacement vector, *E. coli* Amp$^R$, *S. aureus* CM$^R$ | (4) |
| pMK4 | Shuttle vector, *E. coli* Amp$^R$, *S. aureus* CM$^R$ | (5) |
| pXen-1 | Luciferase reporter plasmid, *E. coli* Amp$^R$, *S. aureus* CM$^R$ | (6) |
| pJB67 | pCN51 with optimized RBS and Cd-inducible promoter, *E. coli* Amp$^R$, *S. aureus* Ery$^R$ | (7) |
| pICS3 | RNA expression vector, *E. coli* Amp$^R$, *S. aureus* CM$^R$ | (8) |
| pCN33 | mRNA target GFP reporter, *E. coli* Amp$^R$, *S. aureus* Ery$^R$ | (8) |

| | | |
|---|---|---|
| pGEM-T | pGEM-T Easy Vector cloning plasmid | Promega pGEM-T Easy Vector Kit |

**SSR42 mutant strain construction:** All primers used in this study are listed in Supplemental **Table S4**. A full deletion of SSR42 from the *S. aureus* chromosome was created using the pJB38 allelic replacement vector as previously described (4). Approximately 500-nt upstream and downstream of SSR42 were amplified using primer pairs OL2939/OL3592 and OL3593/OL2389, respectively. These fragments were ligated into pJB38 by restriction digest using enzymes MluI, KpnI, and EcoRI (New England Biolabs). Ligations were transformed into chemically competent *E. coli* DH5α, selected for on LB agar supplemented with ampicillin, and plasmids were purified via Miniprep kit (Qiagen). Correct clones were PCR-confirmed and sequenced using primers OL2380/OL2381, resulting in plasmid pJB38::ΔSSR42. This plasmid was then electroporated into restriction-deficient *S. aureus* RN4220 and selected for on TSA supplemented with chloramphenicol at 30°C. Using phi11 transduction, pJB38::ΔSSR42 was transferred into the USA300 wild-type at 30°C followed by growth at 42°C with chloramphenicol to promote chromosomal integration of the plasmid. Finally, recombinants were grown on TSA supplemented with anhydrotetracycline to select for plasmid excision. Deletion of SSR42 was screened for by PCR using primer pair OL2939/OL2389 and confirmed via sequencing, resulting in the ΔSSR42 strain. Confirmation of SSR42 deletion was validated using RNA sequencing.

**Supplemental Table S4.** Primers used in this study.

| Primer | Sequence* | Description† |
|---|---|---|
| OL2939 | ATGGGTACCCGTTATCTTGTTGGAAGT | F1 SSR42 deletion |
| OL3592 | TCAGCAATTCATACGCGTGCACCAAATAATTT AATTAGACTC | R1 SSR42 deletion |
| OL2389 | ATGGAATTCAATCGCTGTAACGGATTCAT | F2 SSR42 deletion |
| OL3593 | ATTATTTGGTGCACGCGTATGAATTGCTGAAC TCCAATG | R2 SSR42 deletion |
| OL2380 | GCCACCTGACGTCTAAGAAACC | F MCS pJB38ΔSSR42 |
| OL2381 | GCGATTTACATATGAGTTATGC | R MCS pJB38ΔSSR42 |
| OL2888 | ACTGGATCCGTGTCTATAATACTTTGACCT | F SSR42 complementation |
| OL3734 | ATGGTCGACGTTCAATATTTACTACAAAGTCG TG | R SSR42 complementation |
| OL2393 | TCGTATGTTGTGTGGAATTG | F MCS pMK4 |
| OL2394 | GTGCTGCAAGGCGATTAAG | R MCS pMK4 |
| OL398 | TCCTACGGGAGGCAGCAG T | F 16S |
| OL399 | GGACTACCAGGGTATCTA ATCCTGTT | R 16S |
| OL7625 | CAAATAGCTACTCCAAAACGATTAGTTATAA | F RT-qPCR *lukA* |
| OL7506 | CAGGGTTTTCTACAGTAGCAATTC | R RT-qPCR *lukA* |
| OL2123 | GAG TTG TTA TCA ATG GTC AC | F RT-qPCR *sarA* |
| OL2124 | ACT GCT TTA ACA ACT TGT GG | R RT-qPCR *sarA* |

| | | |
|---|---|---|
| OL2487 | TAGTGGTCCATCAACAG | F RT-qPCR *lukS*-PV |
| OL2488 | ACGTTCTACTTCACTGATA | R RT-qPCR *lukS*-PV |
| OL4036 | GGCTCTATGAAAGCAGCAGATA | F RT-qPCR *hla* |
| OL4037 | CTGTAGCGAAGTCTGGTGAAA | R RT-qPCR *hla* |
| OL5048 | GGCAGTGGCTCATTCAACTAC | F RT-qPCR *hlgA* |
| OL5049 | CACCTTTAGAGTTCTGACTTTCTAC | R RT-qPCR *hlgA* |
| OL2170 | GGC TAA AGC TGA ACA TAA TG | F RT-qPCR *splE* |
| OL2171 | ATA TTC ACC ATT GGG ATG C | R RT-qPCR *splE* |
| OL6159 | GTCGCACATTCACAAGTTTATC | F RT-qPCR *aur* |
| OL6160 | GCCTGACTGGTCCTTATATTC | R RT-qPCR *aur* |
| OL4526 | ATGGAATTCGTTGGCATGTCATATTTCTCTCTC CTG | F SSR42 promoter for pXen-1 |
| OL7767 | ATGGGATCCCGAATATTTTGTAACAGGGCTAC TAAG | R SSR42 promoter for pXen-1 |
| OL6607 | AATTGAATTCATTAACTGTCTAGTATCAACGAT C | F *lukAB* promoter for pXen-1 |
| OL6608 | AATTGGATCCAAACACGTTTTTTATTTTTCAT | R *lukAB* promoter for pXen-1 |
| OL7693 | CACGCGCGCAATACTACAG | *perR* screening |
| OL7694 | CGCTTACTTATATGAAGTAAGCTAACG | *perR* screening |

| | | |
|---|---|---|
| OL7501 | AATT**TAATACGACTCACTATAGGG**TCGGATATGCTTAATTTTATAGTAAATTGTATG | F *lukA* 5’ UTR fragment + T7 promoter |
| OL7502 | AAATTGCACATGATAATGATGACG | R *lukA* 5’ UTR fragment |
| OL 7244 | ACTCTGCAGGTGTCTATAATACTTTGACCT | F SSR42 for pICS3 |
| OL 7245 | ATGGAATTCGTTCAATATTTACTACAAAGTCGTG | R SSR42 for pICS3 |
| OL 7341 | AATTAGATCTGTACATTTAATGTTTTTTGATTCAAAC | R SSR42$^{+141}$ |
| OL 7342 | AATTAGATCTGAAGTGATTTCAATCGTACTAATTTTCAGC | R SSR42$^{+230}$ |
| OL 7343 | AATTAGATCTTTGATGTTTTCTAAACAGAACTTTTAAACGC | R SSR42$^{+370}$ |
| OL 7344 | AATTAGATCTCATTATCTTGATGGTGAATTTCGTTG | R SSR42$^{+493}$ |
| OL 7345 | AATTAGATCTTAGATGTTAATTCTTCGCTGCTTAAG | R SSR42$^{+611}$ |
| OL 7346 | AATAGATCTTCTCTTGATGTAAAATCTAAGATGTTTATGC | R SSR42$^{+648}$ |
| OL 7347 | AATTAGATCTATGTGGATGCCTGTCTTGATG | R SSR42$^{+793}$ |

| | | |
|---|---|---|
| OL 7348 | AATTAGATCTATGTAGAAATTGAGTGTGAAAGTTAATAATAGAT | R SSR42$^{+940}$ |
| OL7999 | *GCGCTCCGTC*TACGAAAGAAGGTTATAACAATG | F *lukAB* mutagenesis |
| OL8000 | *CCTGGACGCG*GTATCGACATGTGAATAATATCAC | R *lukAB* mutagenesis |
| OL7499 | AATT**TAATACGACTCACTATAGGG**GATTTCAAACCTATGTATTTC | F SSR42 +T7 promoter for pGEM-T |
| OL7500 | AATATGATGTTCAATATTTACTACAAAGTCG | R SSR42 for pGEM-T |
| OL7501 | AATTT**AATACGACTCACTATAGGG**TCGGATATGCTTAATTTTATAGTAAATTGTATG | F *lukA* UTR +T7 promoter for pGEM-T |
| OL7502 | AAATTGCACATGATAATGATGACG | R *lukA* UTR for pGEM-T |
| OL7523 | AATTT**AATACGACTCACTATAGGG**GCAGATAATTTAGATAAATAAATCATCCATCCATA | F *splE* UTR +T7 promoter for pGEM-T |
| OL7524 | TACCCTCAACCACTGTTGT | R *splE* UTR for pGEM-T |
| OL6163 | ATGGAATTCATGCACCAAATAATTTAATTAGACTCTATCG | F *rsp* promoter for pXen-1 |
| OL6164 | ATGGGATCCCATGTCATATTTCTCTCTCCTGCC | R *rsp* promoter for pXen-1 |

*Underline indicates restriction enzyme site, bold indicate T7 promoter, italics indicates mutated sequence.

†F = Forward; R = Reverse; MCS = Multiple Cloning Site.

**SSR42 complementing strain construction and mutagenesis:** To construct a SSR42 complementing strain, SSR42$^+$, the SSR42 transcript and its native promoter and terminator were amplified using primers OL2888/OL3734 and cloned into pMK4 using restriction enzymes BamHI and SalI (New England Biolabs) to create pMK4::SSR42. This construct was then electroporated into *S. aureus* RN4220 and transduced into the ΔSSR42 strain via phi11 transduction. Wild-type empty vector (WT) as well as ΔSSR42 empty vector strains were also generated using phage transduction of an empty pMK4 vector. To create complements with truncated lengths of SSR42, fragments of SSR42 were amplified using OL2888 along with OL7316, 7337, 7338 to create SSR42-648 nt, SSR42-793 nt, and SSR42-940 nt, respectively. The BlaZ transcriptional terminator (TT) was amplified from the pJB67 plasmid using primers OL7317/7318. The SSR42 fragments were digested with SalI and BamHI and the *blaZ* TT was digested with SalI and PstI. Following restriction digest, all elements were ligated into pMK4, transformed into *E. coli,* and transduced into ΔSSR42 as described above. The resulting constructs included the full-length complement (SSR42$^+$) as well as all truncated complements (SSR42$^{+xnt}$). All pMK4 complements were confirmed via PCR and sequencing using primers OL2393/2394. Mutagenesis was performed using the NEB Q5 mutagenesis kits using manufacturer instructions. Mutagenesis of *lukAB* was performed using primers OL7999/8000.

**Transcriptomic analysis via RNA sequencing:** Wild-type and ΔSSR42 strains were grown overnight in biological triplicate and standardized to an $OD_{600}$ of 0.05 in 100mL TSB. Next, 5mL samples were harvested after 15h of growth, added to equal amounts of ice-cold PBS, and pelleted by centrifugation at 4°C. Total RNA was isolated from cell pellets using an RNeasy Kit (Qiagen), followed by removal of DNA using a TURBO DNA-free kit (Ambion). DNA removal was confirmed via PCR using OL398/OL399 and sample quality was assessed using an Agilent RNA6000 Nano Kit and an Agilent 2100 Bioanalyzer system. After ensuring all samples had an integrity score of $\geq$ 9.9, biological triplicate samples were pooled at equal RNA concentrations. mRNA enrichment was performed by first removing rRNA using a Ribo-Zero Kit for Gram-Positive bacteria (Illumina) before enrichment using a MICROBExpress Bacterial mRNA enrichment kit (Agilent). Library preparation was performed following a Truseq Stranded mRNA Kit (Illumina), omitting mRNA enrichment steps. Sample quality was assessed as detailed above. To quantify library concentrations for pooling of barcoded samples, RT-qPCR was performed using a KAPA Library Quantification kit (KAPA Biosystems). Samples were run on an Illumina Next seq with 150-cycle NextSeq Mid Output Kit v2.5.

**Bioinformatics for RNA-seq analysis:** Raw data was exported in fastq format from BaseSpace (Illumina) and analyzed using CLC Genomics Workbench 20 (Qiagen Bioinformatics). Failed reads were removed using the Illumina Paired Importer tool with quality score parameter option set to Illumina Pipelines 1.8 and later. Reads corresponding to rRNA were filtered and removed by aligning to known rRNA sequences while remaining reads were aligned to the *S. aureus* TCH1516 NCBI reference genome (NC_010079.1) using the RNA-seq Analysis tool (v0.1) with default parameters. Gene expression values were calculated using the Expression Browser tool (v1.1) and differential expression values were generated using the Differential Expression in Two Groups tool (v1.1). Differential expression is reported as fold-change of expression for

ΔSSR42 relative to wildtype. Library size normalization is automatically performed using the trimmed mean of M values (TMM) method by the Differential Expression in Two Groups tool. Ontology classification of genes was assigned based on the Kyoto Encyclopedia of Genes and Genome (KEGG) (9). RNAseq data from this study is available under the GEO accession #GSE237701.

**RT-qPCR verification of RNA-seq dataset:** To validate differential expression data from RNA-seq, a selection of genes with varied expression levels were selected for analysis via Real-Time Quantitative Reverse Transcription PCR (RT-qPCR). Strains were grown and total RNA was isolated from cell pellets as described for RNA-seq studies. Samples were reverse transcribed using an iScript cDNA synthesis Kit (Biorad) and RT-qPCR was performed using gene-specific primers (**Table 2**) and TB Green Premix Ex Taq (Takara). Expression levels were normalized to that of 16S rRNA and fold change of expression was determined using the $2^{-\Delta\Delta CT}$ method. A student's *t* test was used to determine statistical significance relative to wildtype expression.

**Cell fractionation:** The ΔSSR42 and wildtype strains were synchronized and grown for 15h as described elsewhere, and cell pellets were harvested via centrifugation. To isolate secreted proteins, 10% trichloroacetic acid was added to supernatants and incubated at 4°C for up to 24 hours; before centrifugation to harvest precipitated protein. The original cell pellet was resuspended in phosphate saline buffer containing 100µg/ml lysostaphin, 100 U/ml DNAseI (Thermo Fisher), 25 U/ml RNAseI (Thermo Fisher), and 2x EDTA free protease inhibitor cocktail (Pierce, Thermo Fisher), before being incubated for 30 minutes at 37°C. Following this, silica lysis beads were added, and the sample was lysed via bead beating. Subsequent centrifugation at 17,000x *g* for 10 minutes fractionated the samples further into soluble (supernatant) and insoluble (pellet) fractions. The insoluble fraction was washed with PBS and centrifuged once again at 17,000x*g* for 5 minutes. All fractions: secreted, soluble, and insoluble, were then resuspended in cell lysis buffer with a final concentration of 5% (w/v) SDS, 50mM TEAB pH 8.5.

**Proteomics sample preparation:** Fractionated protein samples were reduced by adding 20mM DTT and incubated at 95°C for 10 minutes. To remove any non-solubilized proteins, the samples were centrifuged at 17,000xg for 10 minutes. Protein samples were then quantified using the Pierce 660nm assay with ionic detergent compatibility reagents (Thermo Fisher Scientific) and standardized to 50µg. Alkylation of the samples was performed using a final concentration of 40mM iodoacetamide and incubated at room temperature away from light for 30 minutes. The reaction was quenched with 12% phosphoric acid before adding 7x sample volumes of S-trap buffer containing 90% methanol and 100mM TEAB. Samples were applied to the S-trap column and centrifuged at 4,000x*g* for 30 seconds. The S-trap was washed thrice with S-trap buffer and the flow-through discarded. Digestion buffer containing 50mM TEAB and a 1:10 ratio of trypsin:sample was added and incubated overnight at 37°C. Following incubation, peptides were first eluted with 50mM TEAB via centrifugation. The next elution was performed using 50mM TEAB with 0.2% formic acid, and the final elution was performed using 50% acetonitrile. Samples were then placed in a vacuum centrifuge where they underwent evaporation until complete dryness. Prior to mass spectrometry analysis, samples were stored at 4°C.

**Mass spectrometry:** Peptides were resuspended in in $H_2O$/0.1% formic acid and separated using a 75 µm x 50 cm C18 reversed-phase high performance liquid chromatography (HPLC) column (Thermo Fisher Scientific) on an Ultimate 3000 UHPLC (Thermo Fisher Scientific) with a 120 minute gradient (2-32% ACN with 0.1% formic acid). Peptides were analyzed on a hybrid quadrupole-Orbitrap instrument (Q Exactive Plus, Thermo Fisher Scientific) and complete survey scans were acquired at a resolution of 70,000. The top 10 most abundant ions were selected for MS/MS analysis.

**Mass spectrometry data analysis:** Raw data files were processed in MaxQuant(10) version 2.0.1.0 and searched against the *S. aureus* USA300 pan proteome (UniProt ID UP000001939). Digestion was set to specific with Trypsin/P and fixed modifications included carbamidomethylation of cysteine residues while variable modifications included oxidation of methionine residues and protein N-terminal acetylation. A 1% protein and peptide false discovery rate was applied when identifying proteins. Perseus software was used for further analysis (11). Using the Pride repository (12), raw data and MaxQuant files were uploaded to the proteomeXchange under the ID PXD043724.

**Construction of luciferase promoter fusions:** To monitor promoter activity, we cloned promoter regions of interest into the luciferase transcriptional reporter plasmid, pXen1 (6). The promoter for *lukAB* was amplified using OL6607/OL6608 and cloned into the plasmid using restriction enzymes BamHI and EcoRI (NEB). The promoter for SSR42 was amplified using OL7763/OL4526 and cloned into pXen-1 using the same restriction enzymes. All fusions were transformed into Dh5α, transduced into *S. aureus* strains and confirmed with primers OL5416 and OL5417 and sequencing.

**Blood survival assays:** Blood survival assays were performed as described previously by our group (13). Briefly, bacterial cultures were grown in biological triplicate overnight before being diluted 1:100 and grown for a further 3h. These synchronized cultures were then used to inoculate 1mL of human, gender-pooled whole blood anticoagulated with sodium heparin (BioIVT) at a final $OD_{600}$ of 0.05. Samples were taken immediately after inoculation (T=0) as well as after 4h of rotation at 37°C. Samples were serially diluted and plated onto TSA to calculate CFU/mL. Data is represented as percent survival in comparison to T=0.

**Construction of RNA and mRNA-GFP reporter plasmids:** SSR42 regulation *in vivo* was monitored via a two-plasmid system optimized for Gram-positive species (8). SSR42 was amplified along with its native promoter and terminator using OL7244/OL7245 and cloned into pICS3 using restriction enzymes EcoRI and PstI. The resulting plasmid (pICS3::SSR42) as well as an empty vector were transduced into the ΔSSR42 strain and confirmed with primers OL7350/OL7351 and sequencing. GFP-translational reporter fusions for the *lukA* mRNA was created by insertion of the 5'UTR and a portion of the coding region into pCN33 under control of a constitutive promoter, in this case the *tufA* promoter from *S. aureus.* The UTR of *lukA* was amplified using OL7428/OL7429 and cloned into pCN33 using restriction enzymes BglII and EcoRV before being confirmed via sequencing. The mRNA reporter was transduced into the ΔSSR42 strain containing the pICS RNA expression plasmids and grown on TSA containing erythromycin (5µg/mL) and chloramphenicol (10µg/mL) to select for both plasmids.

**Luminescent reporter assays with neutrophil exposure:** HL60 cells were differentiated as described above before being seeded into black-walled, clear bottomed 96-well plates at a concentration of 2 x $10^5$ in 100µl of phenol red-free RPMI. Bacterial strains were grown overnight as previously described, washed twice with PBS and standardized to a final $OD_{600}$ of 1.0. Following this, standardized bacterial cells were added to HL-60 cells to reach a final $OD_{600}$ of 0.1. Luminescence and $OD_{600}$ readings were taken immediately after inoculation (T=0) using a Biotek Cytation 5 plate reader, with additional readings taken hourly, between which plates were incubated at 37°C and 5%$CO_2$. Luminescence was normalized to $OD_{600}$ for each well. To validate luminescent reporter data, RNA levels were also quantified using RTqPCR. HL60 cells and bacterial strains were prepared in the same ratio as described above, but volumes were scaled up to a final volume of 40ml of HL60 cells, with experiment performed using T75 tissue culture flasks. After two hours incubation, 200µl samples were withdrawn and luminescence measured as described above to confirm that promoter activity phenotypes were consistent. The remaining volume was pelleted and RNA extracted as described above. RTqPCR was then performed as described above but using OL7965/7966 to quantify *lukAB* transcript abundance in each strain.

**Transcriptional Arrest and qPCR Analysis:** Transcriptional arrest and determination of RNA half-life was performed as described (14). In brief, wild-type, ΔSSR42, and $SSR42^{+}$ strains were grown overnight before being synchronized via a 1:100 dilution in fresh TSB followed by growth for 3h. Cultures were then standardized to $OD_{600}$ 0.05 in 5mLs TSB and grown for 15h. After this time, and prior to addition of rifampin (t=0), 5mLs of culture was collected, immediately combined with ice-cold PBS, and cells were harvested via refrigerated centrifugation. Rifampin was then added to a final concentration of 250µg/mL to each culture. At 5, 10, 15, 30, and 45 minutes posttreatment, 5mLs of culture was collected and immediately combined with ice-cold PBS, and cells were harvested via refrigerated centrifugation. All cell pellets were stored at -80° C. RNA isolation and qPCR were performed as described above for each sample, with quantification of RNA abundance at each timepoint calculated using the 2–ΔΔCT relative to the initial RNA abundance at t = 0. A one phase decay curve was generated using Graphpad Prism to determine the decay rate *k,* which was subsequently used to calculate the RNA half-life using the equation $t_{1/2}$ = ln(2)/k.

**References**

1. Salisbury V, Hedges RW, Datta N. 1972. Two Modes of "curing" transmissible bacterial plasmids. Journal of General Microbiologyu 2:443–452.

2. Kolar SL, Nagarajan V, Oszmiana A, Rivera FE, Miller HK, Davenport JE, Riordan JT, Potempa J, Barber DS, Koziel J, Elasri MO, Shaw LN. 2011. NsaRS is a cell-envelope-stress-sensing two-component system of Staphylococcus aureus. Microbiology (N Y) 157:2206–2219.

3. Fey PD, Endres JL, Yajjala VK, Widhelm TJ, Boissy RJ, Bose JL, Bayles KW. 2013. A genetic resource for rapid and comprehensive phenotype screening of nonessential Staphylococcus aureus genes. mBio 4.

4. Bose JL, Fey PD, Bayles KW. 2013. Genetic tools to enhance the study of gene function and regulation in Staphylococcus aureus. Appl Environ Microbiol 79:2218–2224.

5. Sullivan MA, Yasbin RE, Young FE. 1984. New shuttle vectors for Bacillus subtilis and Escherichia coli which allow rapid detection of inserted fragments. Gene 29:21–26.

6. Francis KP, Joh D, Bellinger-Kawahara C, Hawkinson MJ, Purchio TF, Contag PR. 2000. Monitoring bioluminescent Staphylococcus aureus infections in living mice using a novel luxABCDE construct. Infect Immun 68:3594–3600.

7. Windham IH, Chaudhari SS, Bose JL, Thomas VC, Bayles KW. 2016. SrrAB modulates Staphylococcus aureus cell death through regulation of cidABC transcription. J Bacteriol 198:1114–1122.

8. Ivain L, Bordeau V, Eyraud A, Hallier M, Dreano S, Tattevin P, Felden B, Chabelskaya S. 2017. An in vivo reporter assay for sRNA-directed gene control in Gram-positive bacteria: Identifying a novel sRNA target in Staphylococcus aureus. Nucleic Acids Res 45:4994–5007.

9. Kanehisa M, Sato Y, Kawashima M, Furumichi M, Tanabe M. 2016. KEGG as a reference resource for gene and protein annotation. Nucleic Acids Res 44:D457–D462.

10. Cox J, Mann M. 2008. MaxQuant enables high peptide identification rates, individualized p.p.b.-range mass accuracies and proteome-wide protein quantification. Nat Biotechnol 26:1367–1372.

11. Tyanova S, Temu T, Cox J. 2016. The MaxQuant computational platform for mass spectrometry-based shotgun proteomics. Nat Protoc 11:2301–2319.

12. Perez-Riverol Y, Csordas A, Bai J, Bernal-Llinares M, Hewapathirana S, Kundu DJ, Inuganti A, Griss J, Mayer G, Eisenacher M, Pérez E, Uszkoreit J, Pfeuffer J, Sachsenberg T, Yilmaz Ş, Tiwary S, Cox J, Audain E, Walzer M, Jarnuczak AF, Ternent T, Brazma A, Vizcaíno JA. 2019. The PRIDE database and related tools and resources in 2019: Improving support for quantification data. Nucleic Acids Res 47:D442–D450.

13. Brittney D. Gimza SMM& LNS. 2021. An Ex Vivo Model for Assessing Growth and Survivability of Staphylococcus aureus in Whole Human Blood. Methods in Molecular Biology 2341:127–131.

14. Tomlinson BR, Denham GA, Torres NJ, Brzozowski RS, Allen JL, Jackson JK, Eswara PJ, Shaw LN. 2022. Assessing the Role of Cold-Shock Protein C: a Novel

Regulator of Acinetobacter baumannii Biofilm Formation and Virulence. Infect Immun 90:1–21.

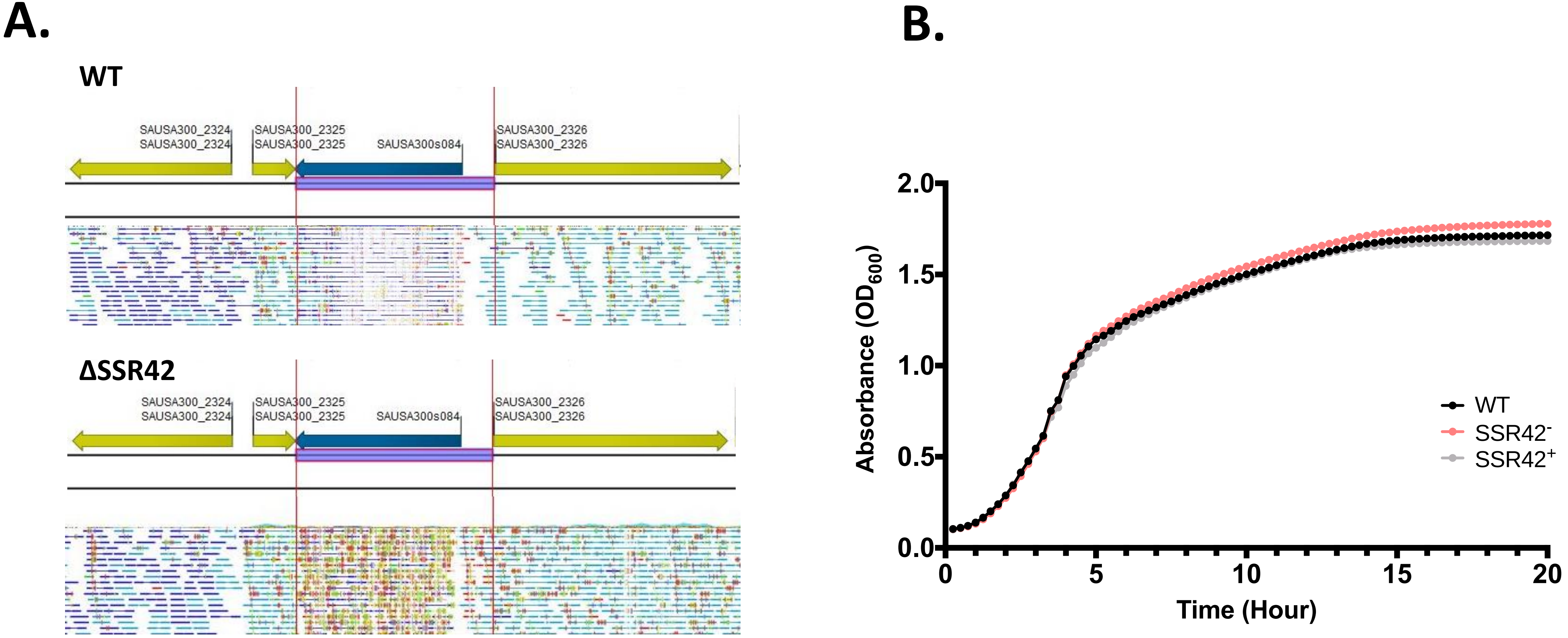


**Figure S1. Deletion of SSR42 has no deleterious effects on surrounding genes and exhibits wild type-like growth. (A)** RNA sequencing reads in the WT or ΔSSR42 aligned and visualized using CLC Genomics Workbench 20 (Qiagen Bioinformatics). Paired reads are in light and dark blue (indicating gene directionality), non-specific matches are yellow, unaligned ends in light green, broken pair reads in dark red. **(B)** Cells were grown in TSB and assessed by measuring optical density ($OD_{600}$) every 15 minutes. Error bars representing ±SEM are shown, although limited variance is detectable between replicates.

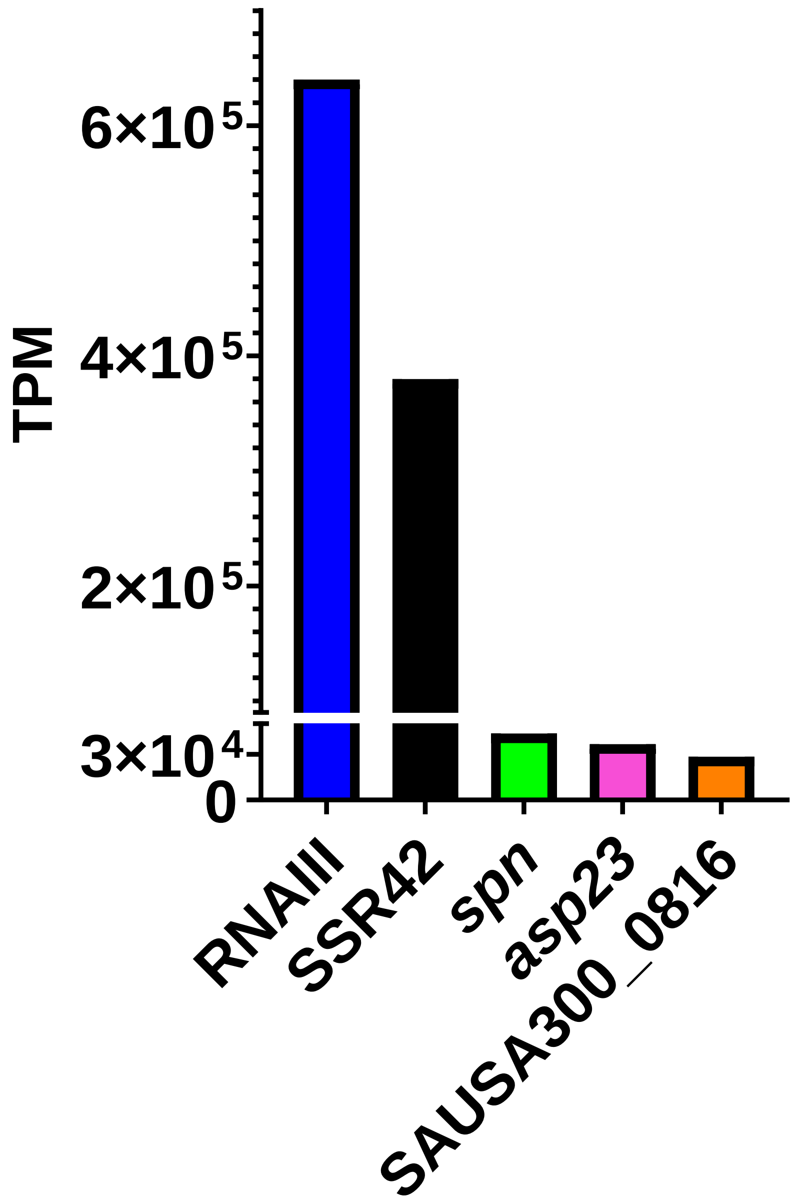


**Figure S2: SSR42 is the second-most abundant RNA in the stationary phase transcriptome.** Wild-type cultures were grown for 15h before RNA was isolated and used for RNA sequencing. Shown are the top five most abundant transcripts from that dataset. Transcript abundance is quantified as transcripts per kilobase million (TPM).

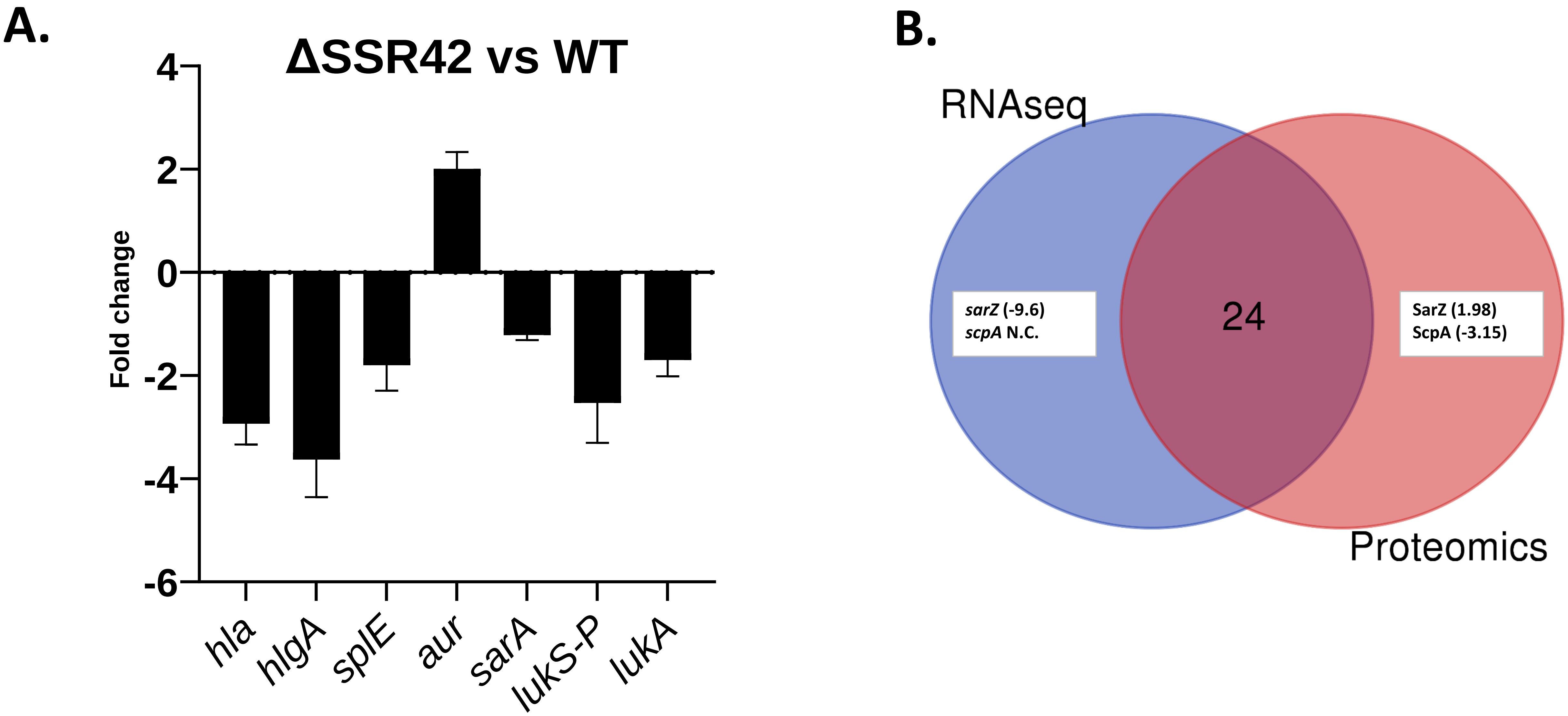


**Figure S3: Validation of RNA-seq findings and comparison to proteomic technologies. (A)** RT-qPCR was performed to assess gene expression between the *SSR42* mutant strain relative to wildtype after 15 h of growth in TSB. The 16S rRNA gene was used as an internal control. Fold change of expression was determined using the $2^{-\Delta\Delta CT}$ method. **(B)** Comparison between RNAseq and proteomics data. Of the 26 key virulence-related factors selected, all trends correlated between methods except for the two labeled. Shown are their corresponding fold changes determined via each method. More detail can be found in Table S1.

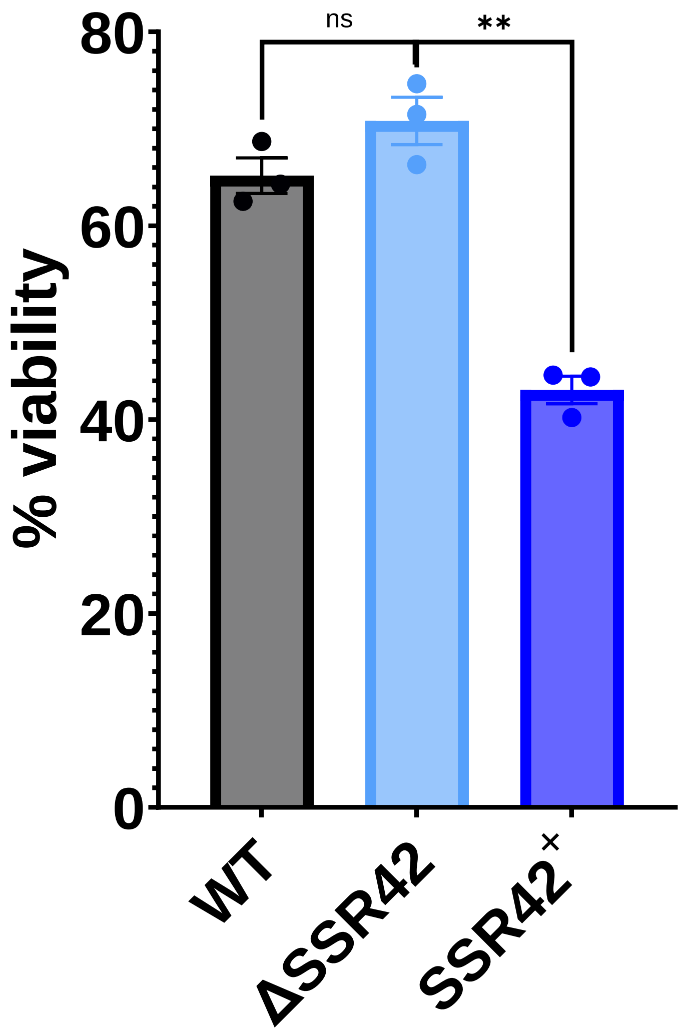


**Figure S4: Neutrophil viability post infection.** As in Figure 2C however viability of the neutrophil-like cells was determined after 24h using the CellTiter reagent. Percent cell viability in comparison to uninfected cells is shown. All experiments were performed in biological triplicate. Error bars represent ±SEM. Student's t test with Welch's correction was used to determine statistical significance. ns – not significant, ** P≤0.01 .

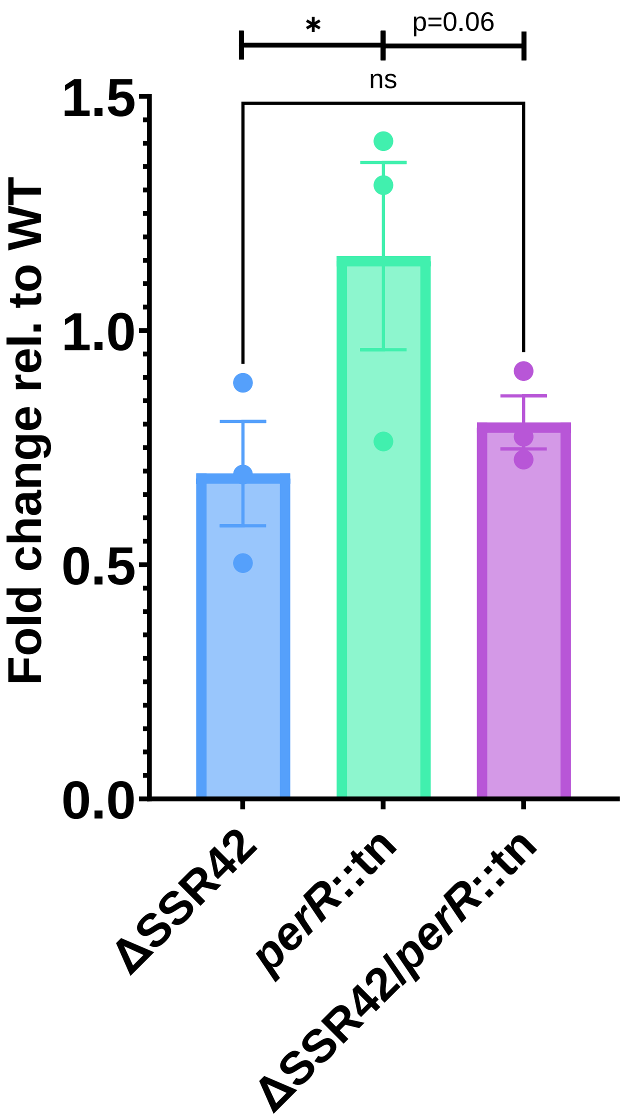


**Figure S5: SSR42 controls LukAB expression in response to neutrophils.** Differentiated HL60 cells were seeded at a concentration of 2x10[5] before the WT and strains shown were added at a final $OD_{600}$ of 0.1. Samples were exposed for 2hrs, pelleted via centrifugation, and RNA extracted. qPCR analysis was then performed for *lukAB* transcript abundance, which was calculated using the $2^{-\Delta\Delta CT}$ method and represented as fold change relative to WT. Data is from three biological replicates. A student's t test with Welch's correction was used to determine significance. Error bars are ±SEM. *$P \leq 0.05$.

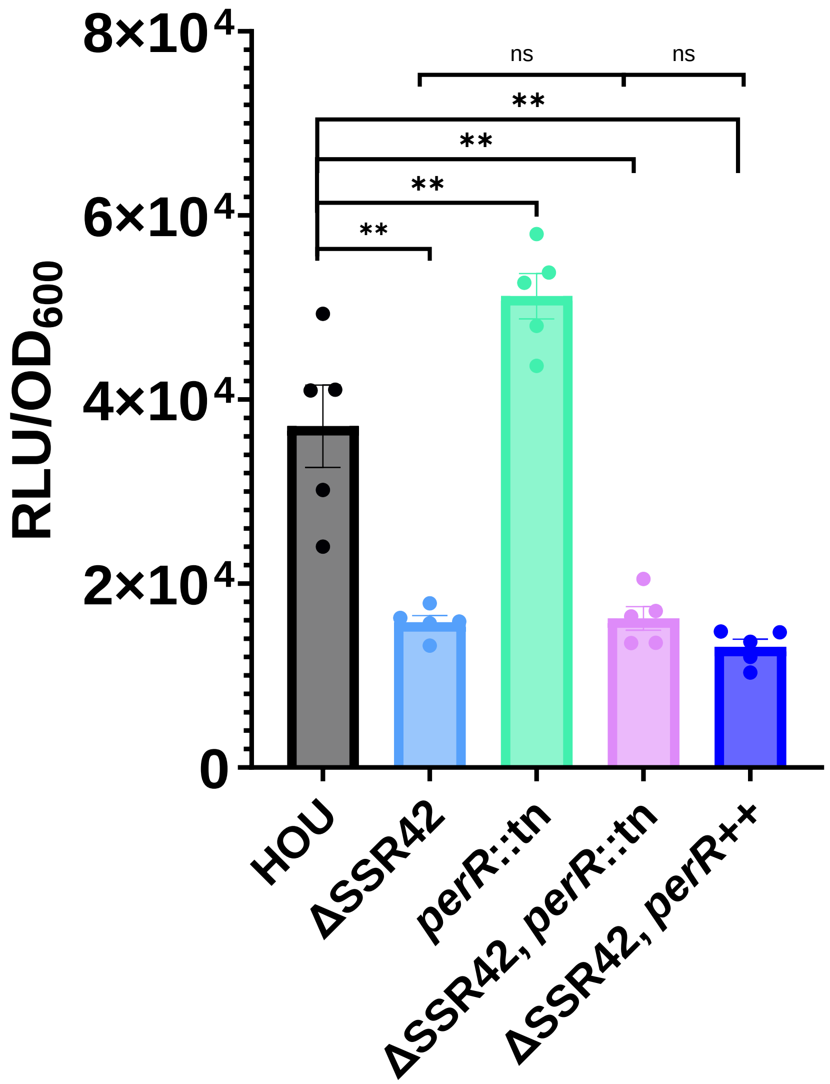


**Figure S6: SSR42 mediates *lukAB* induction independently of PerR overexpression.** As in Figure 3A but using the *S. aureus* strains shown, all harboring a $P_{lukAB}$-*lux* reporter fusion. Luminescence and $OD_{600}$ reads were taken after 2 hours. Luminescence was normalized to $OD_{600}$. For overexpression of *perR* (*perR*++) the pJB67 plasmid was used and cadmium chloride added at a final concentration of 40µM/mL for induction. Data is from five biological replicates. A student's t test with Welch's correction was used to determine significance. Error bars are ±SEM. ** $P \leq 0.01$.

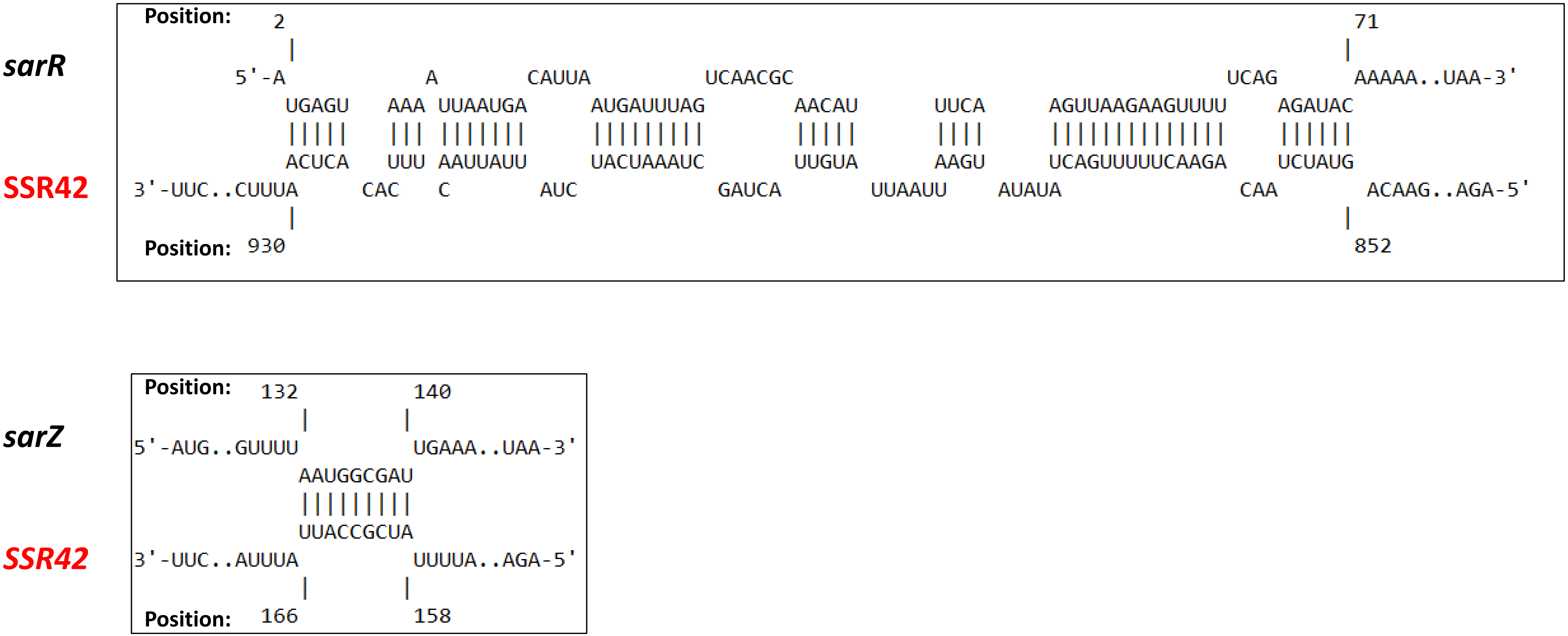


**Figure S7: Predicted RNA-RNA interactions between SSR42 and SarA-homolog mRNAs** IntaRNA was used to identify thermodynamically likely regions of base paring between SSR42 and the *sarR* and *sarZ* mRNAs.

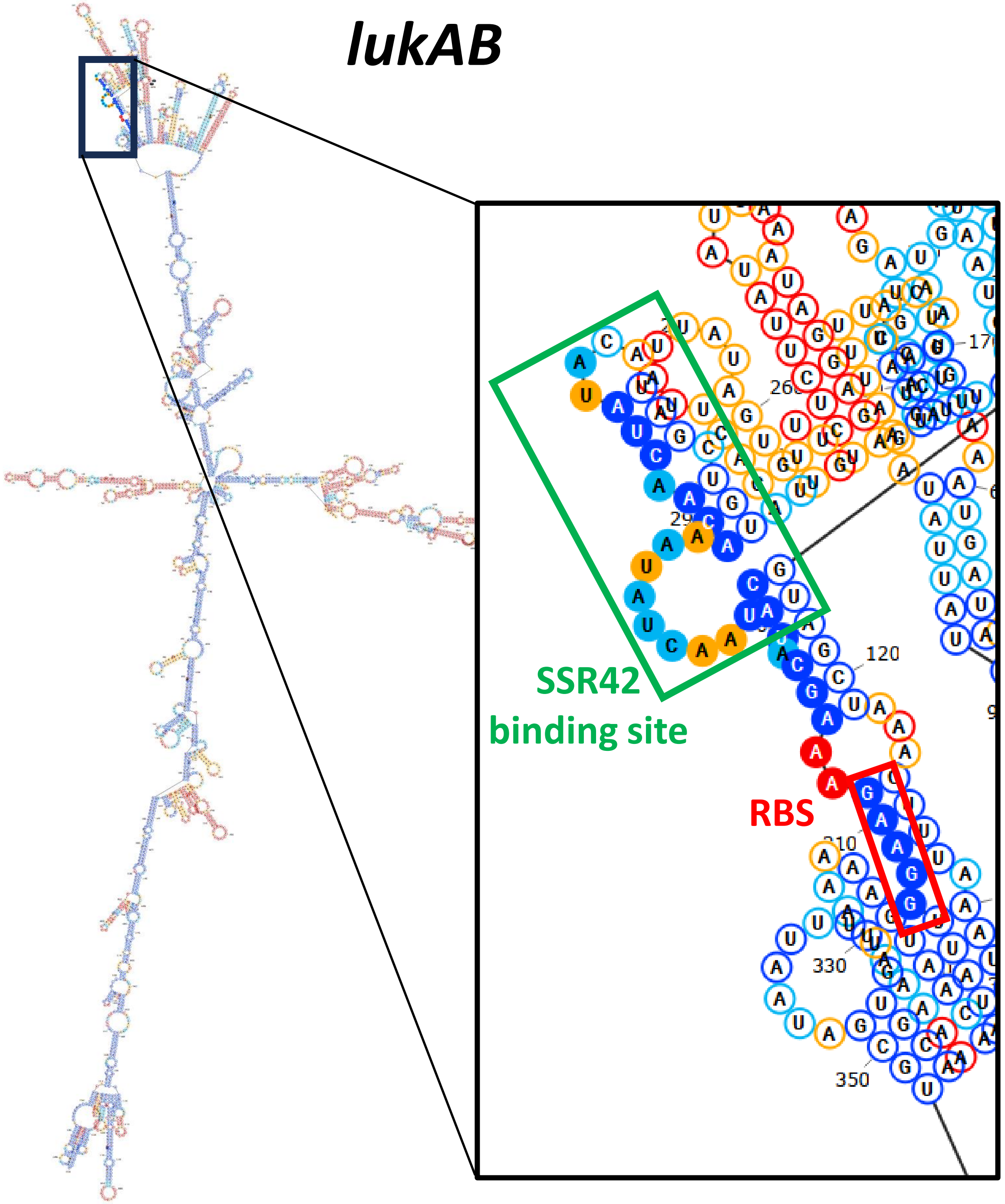


**Figure S8: Predicted secondary structure of *lukAB* mRNA.** The *lukAB* transcript is shown with features of interest labeled. Secondary structures predicted using the ViennaRNA package and visualized using Snapgene.

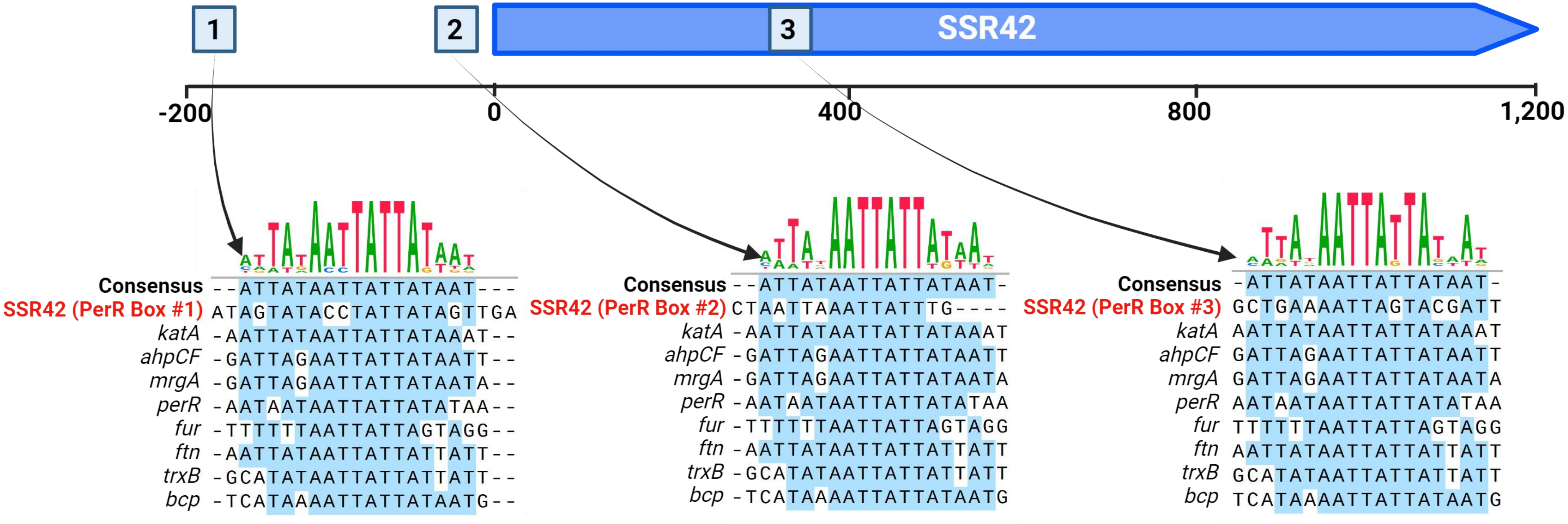


**Figure S9: Identification of PerR binding boxes within the SSR42 promoter region.** Sequences were aligned using MUSCLE to known PerR boxes and consensus sequence. Putative PerR binding regions are highlighted. Figure created using Biorender.com.

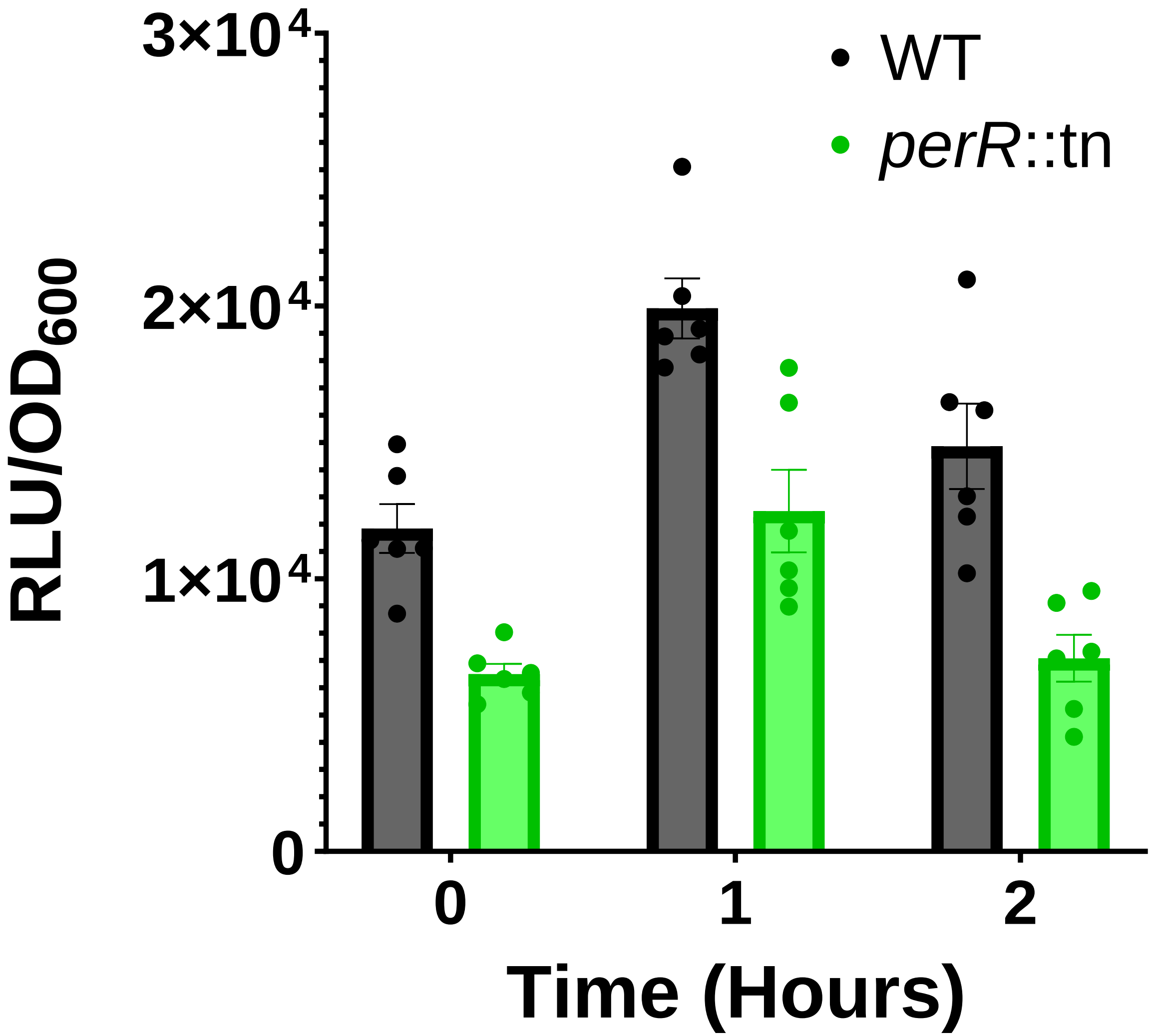


**Figure S10:P$_{rsp}$ does not undergo derepression via PerR in response to neutrophils.** Differentiated HL60 cells were seeded at a concentration of 2x10$^{5}$ before *S. aureus* strains harboring a P$_{rsp}$-*lux* reporter fusion were added at a final $OD_{600}$ of 0.1. Luminescence and $OD_{600}$ reads were taken after 2 hours. Luminescence was normalized to $OD_{600}$. Data is from six biological replicates. Error bars represent ±SEM
